# Modeling the dynamics of within-host viral infection and evolution predicts quasispecies distributions and phase boundaries separating distinct classes of infections

**DOI:** 10.1101/2021.12.16.473030

**Authors:** Greyson R. Lewis, Wallace F. Marshall, Barbara A. Jones

## Abstract

We use computational modeling to study within-host viral infection and evolution. In our model, viruses exhibit variable binding to cells, with better infection and replication countered by a stronger immune response and a high rate of mutation. By varying host conditions (permissivity to viral entry *T* and immune clearance intensity *A*) for large numbers of cells and viruses, we study the dynamics of how viral populations evolve from initial infection to steady state and obtain a phase diagram of the range of cell and viral responses. We find three distinct replicative strategies corresponding to three physiological classes of viral infections: acute, chronic, and opportunistic. We show similarities between our findings and the behavior of real viral infections such as common flu, hepatitis, and SARS-CoV-2019. The phases associated with the three strategies are separated by a phase transition of primarily first order, in addition to a crossover region. Our simulations also reveal a wide range of physical phenomena, including metastable states, periodicity, and glassy dynamics. Lastly, our results suggest that the resolution of acute viral disease in patients whose immunity cannot be boosted can only be achieved by significant inhibition of viral infection and replication.

**Author summary:** Virus, in particular RNA viruses, often produce offspring with slightly altered genetic composition. This process occurs both across host populations and within a single host over time. Here, we study the interactions of viruses with cells inside a host over time. In our model, the viruses encounter host cell defenses characterized by two parameters: permissivity to viral entry *T* and immune response *A*). The viruses then mutate upon reproduction, eventually resulting in a distribution of related viral types termed a quasi-species distribution. Across varying host conditions (*T, A*), three distinct viral quasi-species types emerge over time, corresponding to three classes of viral infections: acute, chronic and opportunistic. We interpret these results in terms of real viral types such as common flu, hepatitis, and also SARS-CoV-2019. Analysis of viral of viral mutant populations over a wide range of permissivity and immunity, for large numbers of cells and viruses, reveals phase transitions that separate the three classes of viruses, both in the infection-cycle dynamics and at steady state. We believe that such a multiscale approach for the study of within-host viral infections, spanning individual proteins to collections of cells, can provide insight into developing more effective therapies for viral disease.

## Introduction

Viral infections are ubiquitous across the tree of life. Though great progress has been made in the prevention and treatment of diseases caused by viruses, a deep understanding of how they infect hosts and evolve within those hosts remains elusive. Of the many ways in which viruses cause disease and evade a host’s immune response, one common theme is a high rate of mutation during replication, leading to the formation of a cloud of genetically similar viral progeny known as a quasispecies [1–5].

Over the past four decades, a growing body of work has demonstrated experimentally the existence of viral quasispecies in infections for diseases including polio, hepatitis C, SARS-CoV-2, and others [6–13]. Additionally, it was found that different viruses within a quasispecies can exhibit a wide range of infectiousness, virulence, and replicative fitness [6, 14, 15]. Complementary to these developments, the theoretical foundations of quasispecies, proposed originally by Eigen, have been the subject of extensive study in mathematical biology and physics, leading to exact solution methods and applications ranging from B-cell receptor diversity to intra-tumor population dynamics [16–20].

Approaches for understanding within-host viral infections based in theory and computation have grown substantially in both number and complexity [21]. Computational approaches range from agent-based models (e.g. [22]), to simulation of partial differential equations (e.g. [23]), to multi-compartment hybrid models (e.g. [24]), and beyond. Although less common, approaches based on statistical mechanics have previously proven effective and provided insight into both biology and physics [25]. A recent study describes a model linking fitness distributions to the probability of causing a pandemic [26]. Our work is built off of that by Jones et al. [27], in which statistical mechanics and thermodynamics are used to better understand viral quasispecies infections within hosts at steady state.

In this work, we model and study the dynamics of viral quasispecies inside an individual host via a 3-step process (Fig. 1). Using a novel, exact calculation method, we determine the distribution of viral subclasses (“match numbers” *m*) across a finite-sized and self-replenishing pool of infectible host cells that have a limited viral capacity. After infection, the host mounts an immune response, clearing viruses in proportion to their match number. Viruses that survive immune clearance undergo replication with a probability that increases with match number, inducing a population-level tug-of-war via the opposing pressures of replication and the immune response. Viruses that do replicate are subject to the additional pressures of mutation, while those that do not will remain inside cells into the next round of infection, effectively shrinking the available pool of cells that newly produced viruses can infect. We allow only one mutation per virus per iteration, and our mutation matrix has a natural fixed point around *m* = 14, a moderate value.

**Fig 1.**
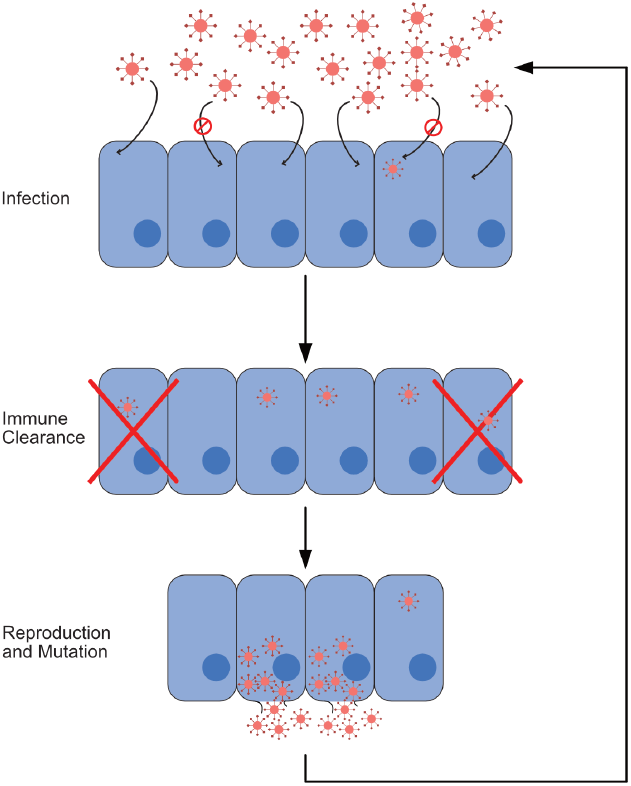
Infection process. In this figure, we show the cycle composing a single iteration of our model. This cycle consists of three steps: infection, immune response, and reproduction with mutation. Viruses that reproduce and mutate from the last step form the cohort infecting cells at the beginning of the next iteration. In our model, one iteration corresponds to a single time step.

To describe a broad variety of host environments, we model host cells as exhibiting a specified level of viral permissivity *T* : low values of T allow only viruses with the highest match numbers to infect, while high values of T permit viruses of all types to infect and replicate. Additionally, based on prior work, *T* can act as a thermodynamic, but not physiological, temperature [27]. At the same time, we model a wide range of host immune responses by subjecting viral infections to different values for the maximal immune clearance (A) of a sigmoidal immunity function.

By varying T, A, and the initial infection, we study infection dynamics across a large space of possible host-virus interactions. The combined pressures of permissivity, immunity, error-prone replication, mutational drift, and the pool of infectible cells’ limited size lead to a phase space with a rich variety of viral behaviors and physical phenomena. We find that the T-A phase space is separated into three main non-extinct regions; and that those regions correspond well to the physiological classes of acute, chronic, and opportunistic infections. We also find that the boundaries in phase space separating the three infection regimes range from first to infinite order, and that infections near the first-order transition exhibit behavior characteristic of “glassy” dynamics. Lastly, we make two predictions: that chronic viral infections are unlikely to be cured through increasing immunity alone, and that the transition from acute to chronic infection behavior is discontinuous across different viral subtypes.

## Model, methods, and materials

Our system consists of a finite pool of cells and viruses that may infect those cells. Viruses can reside either inside cells or in the environment/extracellular space. The infection cycle consists of three main steps: 1) attempted infection of cells by viruses in the environment, 2) suppression of infection by immune clearance, and 3) either viral replication (leading to cell lysis and viral escape into the environment) or latency (persistence inside cells without replication) (Fig. 1). We represent each stage of the infection cycle by the vectors *P* (pre-infection in environment), Ψ^*I*^ (infection), Ψ^Ξ^ (immune response), and Ψ^*R*^ (remaining in cells). After the third step, the cycle begins again, repeating until a number of iterations sufficient to reach steady state is reached in the simulation. We define each iteration as a single time step in the dynamics.

### Model

In order to capture essential information about variation within and across viral quasispecies, we describe each virus in our model with a “match number” *m*. The match number characterizes a cell-virus interaction and can be interpreted as a measure of the binding strength of an idealized viral surface protein to a simple host-cell receptor protein. In our model, all host cells express a single relevant receptor type of sequence length 50 with unchanging identity, while viruses exhibit a single surface protein with varying composition and length 100. The optimal alignment (which may contain gaps) between these two sequences sets the value of *m*, which ranges from 0 (minimal alignment) to 50 (maximal alignment). The match number enables us to group equivalent interactions among a statistically large number (in total, 26^100^) based on an extended amino acid set [27]) of possible viral surface protein compositions.

Viruses in our system are subject to five main pressures. The first two are permissivity and immunity, which directly oppose each other for a given viral match number: a high *m* enables more effective infection and reproduction at the cost of greater immune clearance, while a low *m* enables immune evasion at the cost of decreased infective and reproductive ability. The third pressure is mutation, as random sequence changes to viral offspring can lead even the most infectious virus to have less effective progeny. The fourth, an entropic pressure, is the mutationally-based limitation on variety of offspring that a virus can produce - there are far more potential progeny for some match numbers than others, given a finite alphabet for constructing proteins. The final pressure is induced by the finite pool of cells, as new viruses must compete with both each other and latent intracellular viruses for the limited number of cells to infect. Throughout this work, we see the effects of the interplay between these pressures.

Our model differs from standard quasispecies modeling approaches in four key ways. Firstly, we include a finite and self-replenishing pool of cells in which viruses may remain latent across multiple infection cycles. Secondly, we use a generalized Hamming class (the “match number”) to group viruses with identical numbers of mutations from the optimal sequence, which may not be present at the initial infection. Thirdly, we include mutational backflow, as bidirectional mutation across match number classes is prominent across a large range of *m*. Finally, we perform simulations with large numbers of cells and viruses over long enough timescales to reveal multiple time-dependent features and to ultimately reach steady state.

We now go into more detail about the individual components of the viral life cycle.

### Infection

Infection begins with viruses emerging from the environment and hopping from one cell to the next, trying to infect at each one. At each cell, the virus engages in a wide range of binding poses, testing the interaction of its binding protein at each alignment. If the virus infects, it hops no further, inhibiting other viruses from entering that same cell: in our model we allow a maximum of one virus per cell. However, our model is statistical, and therefore the distribution of viruses in cells after infection consists of two contributions: probability from viral distributions that previously infected the cell, and those that have newly infected. The various types of viruses in the cell thereby undergo competition based on the pressures on the system. Finally, if a virus fails to infect any cells, it leaves the system, either carried to another part of the body where it cannot infect or removed as waste through normal filtration and excretion.

We set the probability that a single virus infects an isolated, unoccupied cell to be *e*_*m*_:

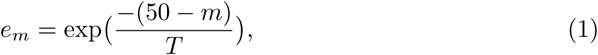

where *m* is the viral match number and *T* is the permissivity. We plot *e*_*m*_ in Fig. 2C.

**Fig 2.**
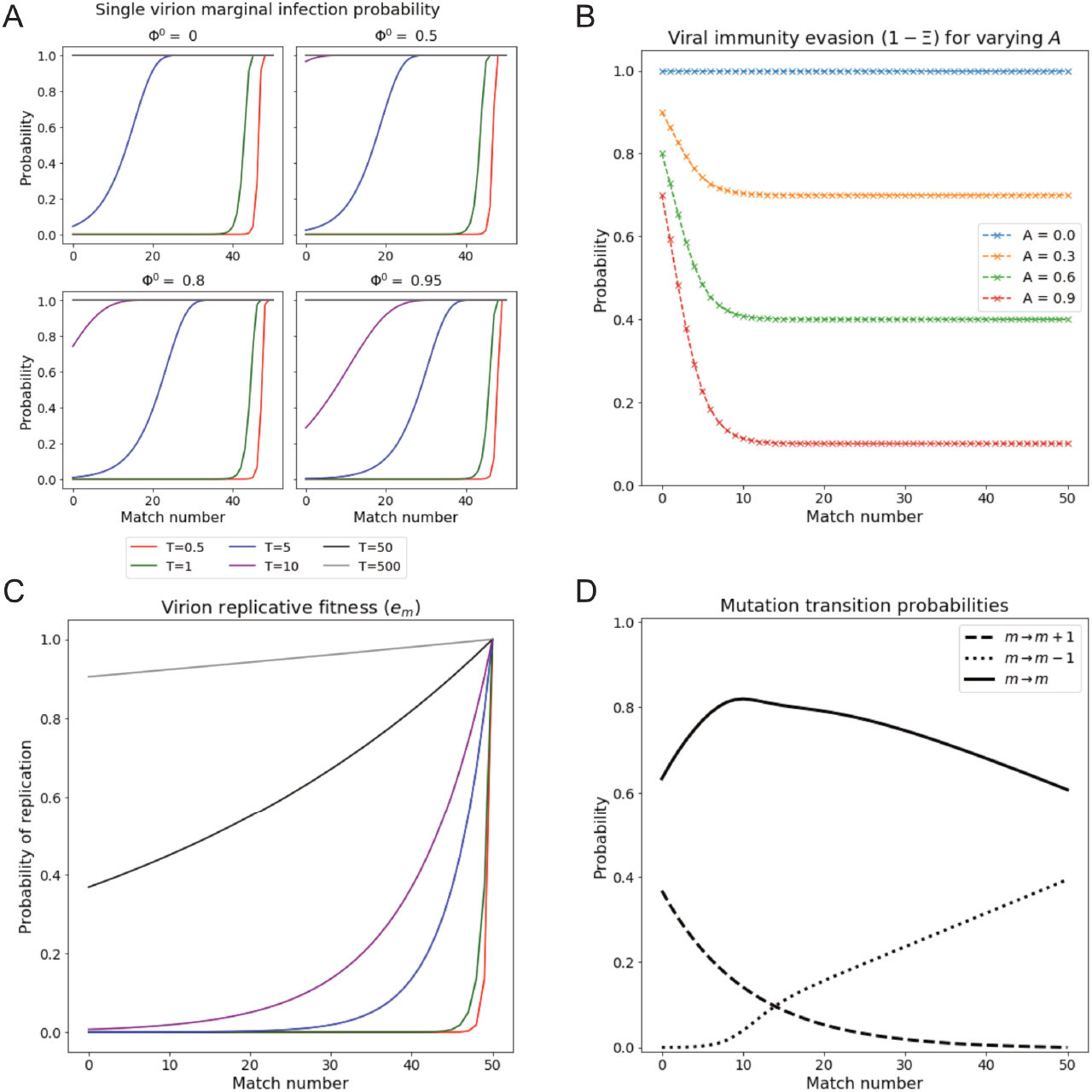
Single virion reproductive fitness and infection probability vary with match number and permissivity. (A) Infection probability of a single virus of match m, under conditions of various prior cell fillings Φ^0^ and for varying permissivity *T*. Higher occupancy results in notable decreases in infection probability especially for lower *m*. Note that infection probabilities, unlike *e*_*m*_ (Panel C), exhibit a shoulder at high *m*. (B) Probability of virus to survive immune response as a function of match m, for various immunity levels *A*. Immune response saturates with increasing *m* above around *m* = 14, where the level is 1 − *A*. (C) Replicative fitness (probability of replication) as a function of match number for varying *T*. Legend shared with Panel A. (D) Replication-associated mutation probability as a function of *m*. Note that the probabilities of increasing and decreasing *m* via mutation are equal at *m* = 14, giving the mode of the Natural initial distribution.

However, the full infection process involves multiple viruses attempting to infect many cells and also takes into account prior cellular infection. Equations governing this behavior are shown in Eqs. 2a-2c, and plotted for a single virus attempting to infect cells of varying prior occupancy level in Fig. 2A (derivation in S1 Appendix, section 1). We note that these equations are an exact representation of the process described above, whereas previous work used approximate expressions to derive analytic results [27].

The result is calculated recursively, so that the infection probability laid down by the first virus, plus the viruses retained after the previous infection cycle, serve as the infected cells that the next virus encounters. For example, the third virus contends with the effects of the first and second viruses, plus those remaining in cells from the previous infection cycle, and so forth, until all *N* viruses at that stage of cycle have had a chance to infect. In order to calculate infection probabilities in a statistical manner consistent with our model, we average over both the order in which viruses encounter cells and the possible sequences in which viruses emerge. We then arrive at the following recursive formulas:

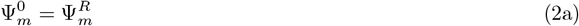

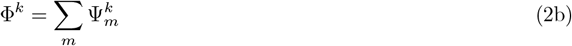

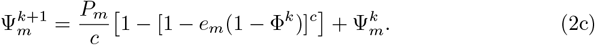

Thus, for a system with *N* viruses, the probability of cells being occupied by viruses after infection is

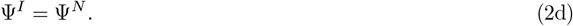

Here, 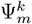 represents the net probability that a single cell will be infected by a virus of match number *m* after the *k*-th virus has tried to infect. As defined above, *P*_*m*_ is the probability distribution over match numbers of viruses in the environment before the infection cycle begins, *c* is the number of cells, and *k* ranges from 0 to *N* (the number of viruses in the environment before the infection cycle begins).

We note that the number of viruses in the environment after replication and mutation is represented by a real number. However, Eq. 2d requires an integer number of viruses. Therefore, to determine the value of Ψ^*N*^ for real *N*, we linearly interpolate the value of Ψ^*N*^ between the two whole numbers of viruses nearest to *N*:

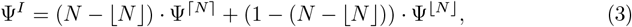

where ⌊*N*⌋ is the greatest integer less than or equal to *N*, and ⌈*N*⌉ is the least integer greater than or equal to *N*.

### Immune response

Our model of the immune response depends only on the match number and the system immune clearance intensity, *A*. The function does not change within a simulation: its static nature assumes that prior infections have created a fixed memory. Those viruses with a higher match number are those that are most likely to have infected in the past and are therefore most likely to be recognized during a new infection. We take the immunity function to have a sigmoidal shape with an inflection point *ν* fixed at *m* = 6 amino acids [27] that sets the 50% immune intensity response:

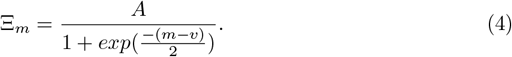

We note that immune clearance is greatest for *m* corresponding to maximal infectious and reproductive ability, setting up a dynamic evolutionary pressure between *T* and *A*. The maximal strength of the immunity we set to *A*, with *A* ranging from zero to one (Fig. 2B). The probability of destroying a virus only approaches 1 with *A* = 1 and moderate match number. Taken together, the viruses that remain after immunity are

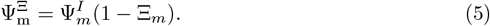

### Replication with mutation

If a virus is able to both infect a cell and evade the immune response, it engages in the third and final stage in its life cycle: replication and mutation. We take the probability to replicate to be *e*_*m*_, the same as the infection probability for a single empty cell, as both replicative and infective ability are essential to a virus’s function.

As depicted in Fig. 2C, viruses with *m* = 50 replicate with probability 1, but for all other match numbers, the probability is less, although at no match number is it exactly zero. However, for any non-infinite permissivity, the probability of replication with very low match numbers is exponentially small. As *e*_*m*_ takes on values from 0 to 1 (with predictable behavior at both extremes), we can derive a natural quantitative scale for permissivity by setting *e*_*m*_ = 0.5 and then determining the upper and lower limits of *T* when *m* = 0 and *m* = 49. This gives a range of *T* ≈ 1 to *T* ≈ 70, where we should expect the majority of interesting behavior in the model to appear.

The distribution of viruses that remain latent or reproduce and escape from cells are, respectively,

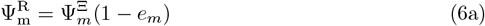

and

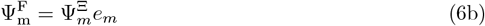

Here, Ψ^*R*^ is the probability that cells remain infected after viruses have a chance to reproduce, and Ψ^*F*^ is the probability that a cell will have had its viruses reproduce. For viruses that do replicate, we set the fecundity (*φ*, number of new viruses produced from each progenitor virus) to 20, giving the total number of viruses present in the environment at the beginning of the next infection cycle:

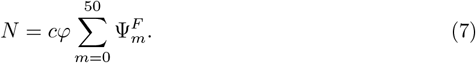

Each viral offspring is mutated at exactly one position in the viral protein sequence [28]. The one amino acid out of 100 on the virus that mutates is chosen at random. This single change can either keep the number of matches to the cellular receptor protein the same, increase it by one, or decrease it by one. Depending on the match number, the probabilities for these three possibilities can shift – large match numbers will have a lower chance for still higher match, and low match numbers a lower chance for still lower match. These transition probabilities were estimated using high-performance computing (HPC) calculations to generate the mutation matrix ℳ (Fig. 2D; described in S1 Appendix, section 2; originally in [27]). The probability distribution of viruses in the environment at the beginning of the next infection process is, then,

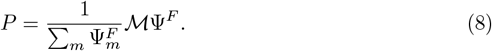

All newly replicated viruses escape from the cell into the extracellular environment for the next cycle. The fixed pool of cells is replenished with uninfected cells to keep the number constant at *c*, and the infection process starts again.

### Computational implementation

Considered across repeated cycles of infection, immune response, and replication, the equations above can be seen as difference equations, which we evaluate through iterative calculation. We run simulations for 100,000 iterations, as this almost always guaranteed convergence in our simulations. We define convergence at the latest iteration *i* such that, for all following iterations,

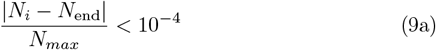

and

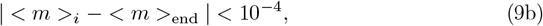

with <*m* >_*i*_ the sample mean of *P*_*m*_ at iteration *i, N*_*i*_ the number of viruses in the environment at iteration *i, N*_*max*_ the maximum allowed number of viruses in the environment (equal to *cφ*), and an “end” subscript refers to the value at the end of a simulation. In addition, given that viral population sizes are real-valued scalars, we set a extinction cutoff of 10^−8^ such that a simulation is terminated when the environmental viral load sinks beneath that cutoff.

For this work, we run simulations across the combination of parameters shown in Table 1.

**Table 1.**
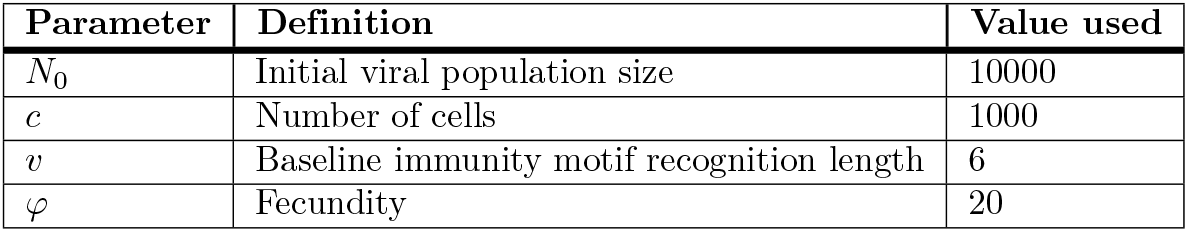
Simulation parameters.

| Parameter | Definition | Value used |
| --- | --- | --- |
| $N_0$ | Initial viral population size | 10000 |
| $c$ | Number of cells | 1000 |
| $v$ | Baseline immunity motif recognition length | 6 |
| $\varphi$ | Fecundity | 20 |

We initialize simulations with either a uniform or “natural” match-number distribution (Figs. 3A, B). The uniform distribution is defined in the standard manner, with each value of *m* assigned equal probability. The “natural” initial distribution is defined as the steady-state distribution for *A* = 0 and *T* → ∞, which is also the stationary eigenvector of the mutation matrix. Using these two distributions enables us to compare effects of the mean and variance of the initial match-number distributions.

**Fig 3.**
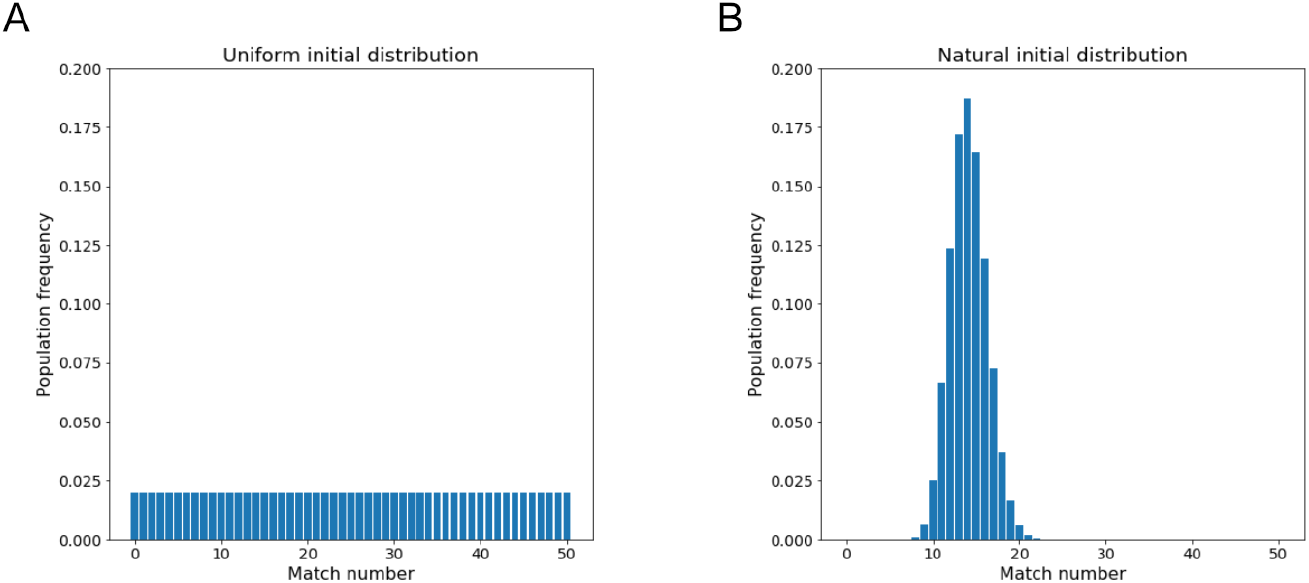
Initial distributions used in simulations. (A) Uniform initial distribution. (B) “Natural” initial distribution (infinite permissivity, zero immunity limiting steady-state distribution).

We also note that in three limits (very low *T*, near infinite *T*, and near-complete cellular occupation after the immune response), we are able to solve for the steady state of the system analytically, suggesting that it is unlikely that there are additional, different steady states that will appear at much later iterations.

Additionally, through roughly approximating the timescale of infection from experimental data, we find that the duration of simulations reaches and may even exceed the timescale of a human lifetime. Therefore, in the unlikely case that the true steady-state is not reached by the end of our simulations, those later states are not relevant to the conclusions of this paper.

We have implemented simulations in Python 3 using Numpy accelerated with Numba package [29, 30]. Where referenced in the code corpus, snippets from Stack Overflow have been used and referenced accordingly. All code is available on the Center for Cellular Construction Github page (https://github.com/cellgeometry/ViralStatMech).

## Results

We first highlight that, independent of initial condition, all viruses in our model coalesce into coherent quasispecies distributions, often after only a few iterations of our simulations. Based on the work of Eigen [1–3], we attribute the emergence of quasispecies distributions to the presence of a mutation matrix in our model. These distributions are nearly always single-peaked, unimodal, and fairly narrow, with a roughly Gaussian shape and a well-defined maximum. However, as discussed below, there are a small subset of cases in which we see two separated peaks arise.

We begin by considering steady-state results.

### Steady-State

#### Probability distributions

In Fig. 4, we plot the end results of simulating the modeling equations above (Eqs. 2a-8) for 100,000 iterations. We see at steady-state that the match-number distributions coalesce into quasispecies distributions, in some places bimodal. We also find that the model goes to the same steady state determined by *T* and *A*, regardless of initial conditions of viral load and match-number distribution (as long as viruses have not gone extinct); this result is not a given for our model, as the infection process is non-linear. In particular, the Natural initial condition leads to a substantially larger extinction region than does the uniform initial condition, suggesting that quasispecies require more viruses at higher match numbers for survival at low permissivity. We note that extinction only seems to occur at low permissivity; at high enough *T*, no value of immunity is able to completely suppress the viral population.

**Fig 4.**
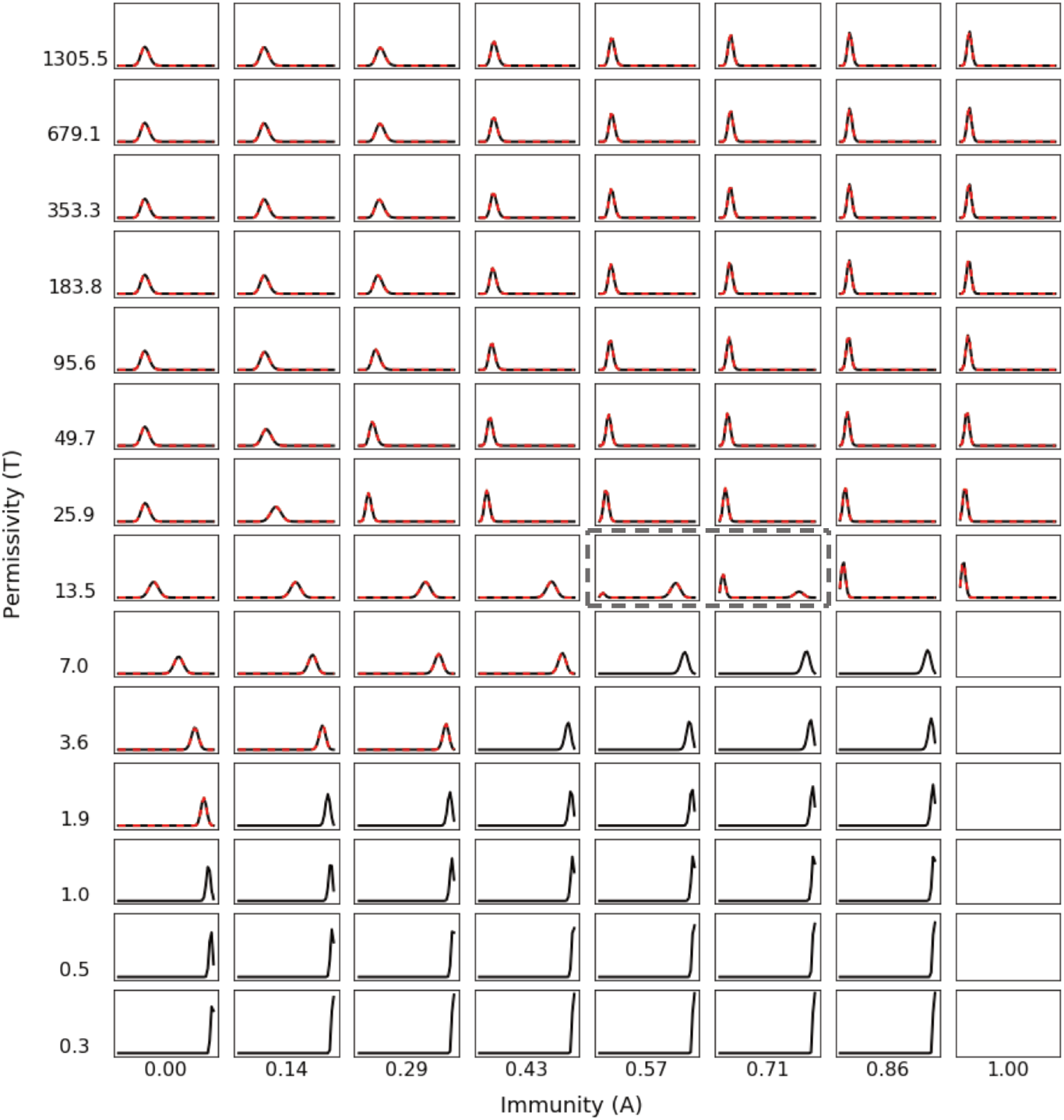
Steady-state match number probability distributions exhibit different regimes of behavior and bimodality near a transition region. Match number distributions at the ends of simulations for a subset of sampled permissivity and immune intensity are shown for uniform (black) and natural (dashed-red) initial distributions. For all simulations, we sample permissivity on a logarithmic scale because *e*_*m*_ is an exponential function of permissivity (Eq. 1). Each distribution is normalized to sum to 1, so broader distribution necessarily exhibit lower peaks. For each distribution, the x-axis ranges in match number *m* from 0-50, and the y-axis ranges ranges in probability from 0-1. The bimodal distributions are enclosed by dark-gray dashed lines and extinction is indicated by the lack of a plotted distribution.

Inspection by eye suggests three general regions of self-consistency. Low permissivity leads to distributions centered around high match numbers, regardless of immunity. High permissivity and moderate-to-high immunity lead to distributions centered around low match numbers. High permissivity and low immunity lead to distributions centered around the natural distribution (3B).

At low permissivity, two effects dominate: the need for viruses to reach very high match numbers in order to both infect and replicate, as well as the effectively *m*-independent suppression of high match-number viruses by the immunity. Accordingly, we observe two trends reflecting these pressures. Firstly, we see that the peaks of the distributions increase toward higher *m* as immunity increases, with the distribution peaks centered at lower *m* for higher permissivity. Secondly, we find that the widths of the distributions decrease as immunity increases and permissivity decreases. These trends are linked and reflect a strategy of the quasispecies to more efficiently increase its population size to overcome increasing immunity. In particular, the distribution peaks shift towards higher *m* and become more narrow in order to take advantage of the sharp increase in replicative ability as match number increases in this permissivity regime (Fig. 2C). Eventually, this strategy fails and viruses are forced into extinction at the highest levels of immunity.

In contrast, in the high-permissivity and higher immunity regime, immunity dominates other pressures in the system. Here, match number distributions are centered at low *m*, tending toward lower *m* as immunity increases. We remind readers that the dynamic range of immunity spans from *m* = 0 to *m* ≈ 15 (Eq. 4 and Fig. 2B). Thus, as immunity increases, viruses are able to evade immune clearance by shifting towards lower and lower *m*. We also see that, as permissivity decreases, the peaks of match number distributions shift to lower *m*, showing the stronger effects of immune pressure over permissivity in this regime. In this regime, permissivity is high enough that viruses with low *m* are still able to infect and reproduce, enabling viruses to pursue both sets of strategies as environmental conditions change. However, as the permissivity further decreases, this strategy becomes untenable as viruses at low *m* become unable to replicate, leading to the emergence of higher *m* viruses at lower permissivity.

Lastly, we address the high-permissivity, low-immunity region. Here, the pressures of permissivity and immunity are minimal, enabling entropic mutational pressures to dominate. Accordingly, we see that match number distributions in this region closely resemble the Natural initial distribution, which is defined as the stationary distribution of the mutation matrix.

We highlight that, broadly across phase space, match number distributions tend to increase in width as immunity decreases. This observation is consistent with previous clinical findings that immuno-compromised or -suppressed patients exhibit greater diversity of viral variants than those with stronger immune systems [13, 31, 32].

We draw attention to the unexpected emergence of bimodal distributions (boxed by gray dashed lines in Fig. 4) at *T* = 13.5 and *A* = 0.57, 0.71. Of note, the peaks of these distributions appear to be centered at the locations of the unimodal peaks at higher and lower *T*.

We also see two major permissivity-dependent trends that span multiple regions. The first, at low immunity, is that distribution peaks shift smoothly to higher match numbers and become more narrow as permissivity decreases. This reflects the steadily growing pressure of permissivity, which increasingly allows only higher match-number viruses to infect and reproduce as *T* decreases. The second, at moderate to high immunity, is that the distribution peaks shift initially towards low *m* and then suddenly jump to increasingly high *m* as permissivity decreases. We can now interpret the emergence of bimodal distributions in the midst of this shift as a point of balance between the pressures of permissivity and immunity.

#### Order parameter

Given the emergence of three distinct regions, a sudden jump in behavior, and the appearance of bimodal distributions, we are prompted to explore the possibility that *T* and *A* define a phase space exhibiting critical behavior. Following prior work on a similar model [27], we define an order parameter:

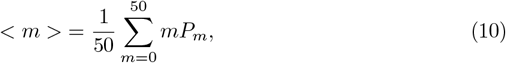

where *P*_*m*_ is the environmental match-number probability distribution at the end of a simulation. As before, we normalize the order parameter to fall between 0 and 1. We note that <*m* > is the sample-mean of the match number distribution for viruses in the environment, generally corresponding to the location of the peak of the associated match number distributions (which are generally symmetric about their centers).

In Fig. 5, we show the the order parameter as a function of permissivity (*T*) and immunity (*A*), revealing a phase diagram with three well-defined phases and varying regions of viral extinction (these results broadly resemble and validate an earlier, approximate form of this model [27]). Using the order parameter, we are able to demonstrate that there are three transition zones: a first-order phase transition, a higher-order phase transition, and a crossover region. A first-order phase transition emerges at moderate-to-high immunity (*A* ≈ 0.2 − 0.9) and *T* ≈ 10 − 20, separating phases I and III. A higher-order continuation of that phase transition separates phases II and III at low immunity (*A* ≈ 0.2 − 0.3) and *T* > 25. Lastly, a crossover region bridges phases I and II at low immunity (*A*< 0.2) and *T* ≈ 10 − 20.

**Fig 5.**
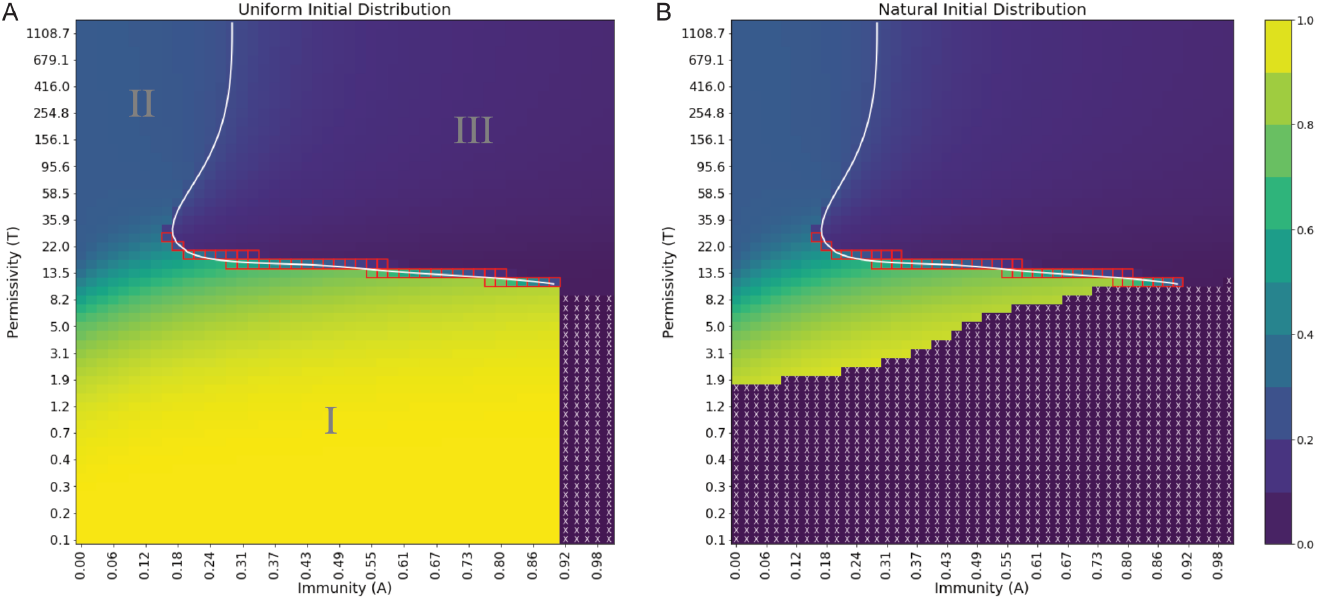
Order parameter at steady state exhibits distinct regions of behavior and extinction, separated by phase boundaries of varying order. (A) Uniform initial distribution. (B) Natural initial distribution. Order parameter values are depicted in a continuous range of colors from purple (0.0) to yellow (1.0). Purple regions with white X’s denote viral extinction before the end of simulations. The white line is a spline fit to locations of maximal difference in order parameter at fixed permissivity or immunity. Red squares denote points where *P*_*m*_ is bimodal. A reminder that the natural scale for permissivity is between *T* ≈ 1 and *T* ≈ 70, where most changes in behavior are expected to occur.

A first-order phase transition can be defined as finite discontinuity in the order parameter (e.g., the change in density upon the melting of ice into liquid water), while a continuous (higher-order) phase transition occurs where the order parameter has a discontinuity in a higher derivative (e.g., a ferromagnetic transition) [33]. A crossover region, also known as an infinite-order phase transition, is a region in which the order parameter of a system undergoes a continuous change between two qualitatively distinct states [34] (e.g., the BEC-BCS transition of ultracold Fermi gases [35]). We note that our model is in the grand canonical ensemble, with an effective bath of “particles” (possible viral sequences) drawn from during replication and returned during infection and the immune response. This bath is therefore of size large enough (26^100^ ≈ 10^140^) to consider the system in the thermodynamic limit, allowing the possibility of phase transitions.

First-order phase transitions in nature often exhibit the coexistence of both states near their transition boundary [33]. We also see this in our results: as shown in Fig. 4, the match number distributions of viruses at the first-order phase boundary are bimodal (enclosed by a dashed-gray box in Fig. 4 and red squares in Fig. 5), with each of the two peaks located at the same position as the unimodal steady-state peaks on either side of the boundary. We interpret this as the coexistence of two different quasispecies.

In addition to the first-order phase transition, there is a second transition that spans moderate-to-high permissivity between low and moderate immunity, which we refer to as the “vertical” phase boundary. This boundary differs in that it changes from second- to higher-order as permissivity increases, smoothly shifting to an infinite-order crossover region as *T* approaches infinity. Even in that regime, the order parameter changes by almost 40% across a region spanning only 0.2 in *A*. Hence, the vertical boundary divides phase space at moderate-to-high *T* into two distinct and separate regions. As described earlier, it marks the boundary between viral match-number distributions that have order parameters more strongly determined by mutational pressure and those determined by immune pressure.

In particular, we note that we do not have a critical point where the phase boundary makes a bend from horizontal to vertical. We do not see a supercritical phase around the cusp, as the crossover region that extends diagonally from that point to *A* = 0, *T* ≈ 5 appears to be continuous for all derivatives. However, we do see some unusual oscillatory behavior in that region for very small *A* around *T* = 5, which we discuss later.

The phase portraits for the two initial conditions exhibit one major difference: in the low-permissivity region (lower half of the phase portrait), we see a dramatically smaller zone of extinction (white X’s) for the uniform initial distribution than for the “natural” initial distribution, which lacks viruses at high *m*.

As noted before, the two phase portraits exhibit identical steady-state results for the order parameters (and indeed, as we shall see below, viral population size) in non-extinct regions. These results, when considered with similar observations for other measurements shown throughout this work, give credence to the existence of a unique non-zero steady-state for each permissivity-immunity pair, a seemingly emergent result that we did not expect given the non-linearity of the model.

#### Viral load

Viral load is, perhaps, the greatest indicator of infection intensity in a clinical context. Here, we explore three types of viral load: intracellular/inside cells, extracellular/environmental, and total (the sum of intracellular and extracellular). Based on the results of this section, we conclude that the three phases correspond to three real disease classes: “acute,” “chronic,” and “opportunistic.” In Fig. 6 we show the steady-state values of these measures for uniform and natural initial distributions. Here, “viral load” is defined as the ratio of the number of viruses at a point in phase space to the maximum allowable viral population (*cφ* = 20, 000 in the environment and *c* = 1, 000 in cells); the total viral load is defined as the sum of these ratios and is the total viral occupation after infection and the immune response. We find that both the intracellular and extracellular viral load landscapes are generally well-partitioned by the order parameter-derived phase boundaries. This observation further reinforces that the phase transitions extend throughout the properties of the system.

**Fig 6.**
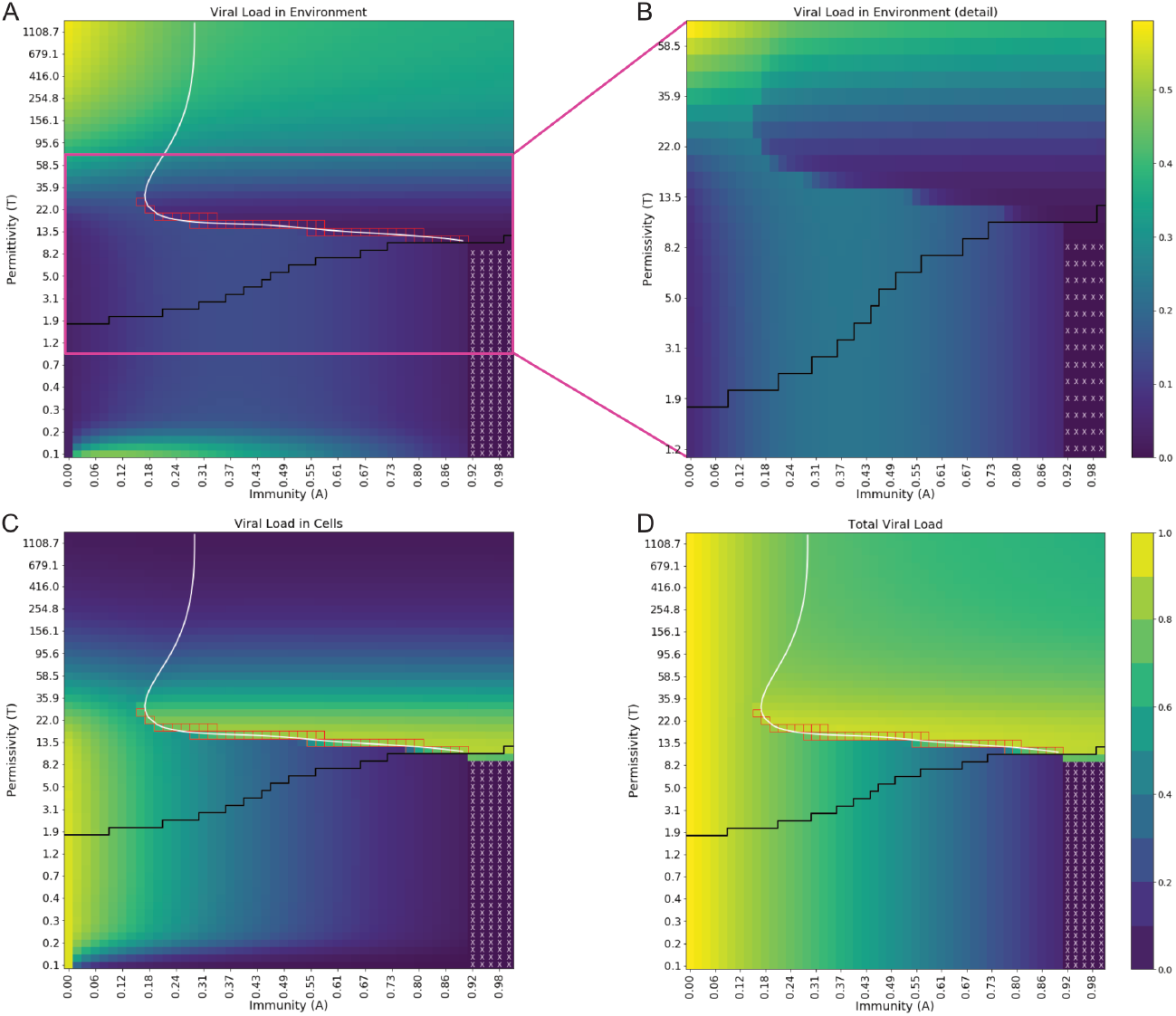
Viral loads in the environment and in cells are well-separated by the phase boundary and demonstrate a fixed-sum rule. (A) Viral load in the environment. (B) Detail of viral load in the environment, confirming discontinuity at the first-order phase boundary. (C) Viral load inside cells. (D) Total viral load. Values shown are from uniform initial distribution simulations but are nearly identical to those from natural initial distribution simulations. Viral loads range from 0 to 1, shown in color ranging from dark purple to bright yellow (0-1, colorbar in D), except for panel (B). White X’s denote points in phase space where uniform initial distribution simulations go to extinction. Points below the black solid line fall in the natural initial distribution extinction regime. White spline is overlaid from Fig. 5, as in other figures. Red squares designate points with bimodal match number distributions at steady state.

Comparing the viral loads in the environment (Fig. 6A) and inside cells (Fig. 6C) shows that, unexpectedly, the total viral load (Fig. 6D) does not depend on either permissivity or immunity for large regions of phase space. This suggests that certain quantities in the model are subject to conservation rules. This effect is most apparent for low immunity (*A*< 0.08), where viruses are predominantly in the environment at high permissivity and gradually shift into cells as permissivity decreases. The largest shift in viral load between cells and the environment takes place roughly around the crossover region (from *T* ≈ 10 to *T* ≈ 20, see Fig. 6B). As permissivity decreases, making infection and replication more difficult, viruses that enter cells are more likely to remain in cells rather than replication at these low levels of immunity. This has two effects: fewer new viruses enter the environment at the end of each iteration, and a dwindling number of cells are available to infect at the beginning of each cycle. Together, these lead to a shift in viral occupation from the environment into cells.

In contrast, we see in phase I that the total viral load depends heavily on immunity. In fact, there is a linear dependence of total steady-state viral load on immunity, as we show analytically in the supplement (S1 Appendix, section 3). Unexpectedly, at very low permissivity (*T* < 0.25) for the uniform initial distribution, the environmental viral load (Fig. 6A) jumps from low values at *A* = 0 to high values at *A* ≈ 0.02, then decreases as immunity increases. To understand this further, at the very lowest value of *T* = 0.1, we are able to use perturbation theory techniques to analytically predict the viral loads: we predict a maximum in viral load that depends on permissivity and find that the results from perturbation theory at *T* = 0.1 match the simulations very well (error < 1%). We include a derivation and further details in the appendix (S1 Appendix, section 4; Supplementary Figs. S1 Fig, S2 Fig).

From these calculations we learn that the change in viral load at low *T* as immunity increases is the combined result of four pressures: the finite number of infectible cells, low permissivity, mutation, and immunity.

We explain these results semi-quantitatively here. At *A* = 0 and very low *T*, the system only allows viruses with either *m* = 49 and *m* = 50 to infect and replicate. At such low *T*, there is a large asymmetry between the infective and reproductive abilities of the two types of viruses: both are able to infect well (Fig. 1A), but *m* = 50 viruses can replicate and escape into the environment far better than *m* = 49 viruses (Fig. 1C). At *A* = 0, both *m* = 49 and *m* = 50 viruses enter cells, but only *m* = 50 leave. When *m* = 50 viruses replicate, they produce a large number of both *m* = 49 and *m* = 50 viruses. Each iteration, additional *m* = 49 viruses enter the cells but do not leave. Eventually, all of the cells are filled with *m* = 49 viruses, preventing *m* = 50 viruses from infecting. The population in cells is maximum and the population in the environment is 0.

As *A* increases, an increasing number of viruses inside cells are killed off. Initially, for low *A*, this allows more *m* = 50 viruses to infect at the beginning of the next infection cycle. However, as *A* increases, this process reaches a maximum, with larger *A* killing off viruses faster than they are able to refill cells, leading the steady-state viral load both inside cells and in the environment to decrease. This trend of environmental viral load growing and then shrinking as *A* increases extends to higher values of permissivity (Fig. 6A) throughout phase I.

In phase III we also see a broad region with high viral load, but in this case it is for the viral load inside cells, and as a function of permissivity rather than immunity. We refer specifically to the region above the horizontal phase boundary, roughly bounded by *T* = 11 and *T* = 20 and *A*> 0.2. We explain this in the following way. Starting from high permissivity, as *T* is lowered at fixed *A*, low-*m* viruses gradually lose the ability to replicate but maintain their ability to infect (Figs. 2A, C). This leads to large accumulation of viruses inside cells, and with low *m* they evade the immune response, resulting in a broad maximum across all *A*. Decreasing *T* still further across the phase boundary, however, enables high *m* viruses to reproduce in sufficient number to overcome the immune response, leading first to coexistence with low *m* viruses around the horizontal phase transition and then outcompeting low *m* viruses into extinction.

Finally, we focus on the high permissivity regime (Phase II, *T* > 130), where at zero immunity the environmental viral load reaches its maximum, as all viruses that infect are able to replicate. As immunity increases, the environmental viral load initially decreases linearly, but the high level of permissivity quickly enables viruses with low match number to survive and replicate regardless of immunity. The viral load in cells is very low in this region because all viruses are likely to replicate, regardless of *m*.

In summary, our analysis of the steady-state results has led to the following findings. First, there are three distinct phases of viral types, separated by phase transitions of varying order and crossover regions. Secondly, the phase transitions and crossover regions seen in the order parameter emerge in the population size landscapes as well. Thirdly, the size and location of extinction regions varies with the initial conditions. Lastly, viral extinction is more likely to occur at lower permissivity and higher immunity.

These results lead us to postulate that each of the three phases corresponds to a disease type affecting humans in real life. Firstly, we suggest that the class of viruses below the horizontal phase boundary (Phase I), due to their persistent reproductive ability at nearly all immunities, corresponds to “acute” viral infections brought on by viruses that are highly adaptable (via mutations) and generally infectious, such as the influenza viruses, rhinoviruses, and others, including SARS-CoV-19 [36, 37]. Also, in this region, the total viral load exhibits a strong inverse dependence on immunity, concluding with regions of total viral extinction for high immunity. Notably, the region of extinction is larger in the case of the more lifelike natural distribution, which does not contain contributions from the extremes of the match number spectrum.

Secondly, we see in Phase III, without the pressure from low permissivity, viruses with low steady-state match number are able to survive while consistently evading the immune response. Even if immunity is increased in this phase, we find that there is no clear resolution of viral infection, consistent with the physiology of long-term, chronic diseases. We thus associate this phase with “chronic” viral infections, as this behavior matches well onto the set of viruses that stay primed in the body for many years, such as varicella-zoster (chicken pox/shingles), hepatitis B and C, and HSV [38–41].

Lastly, in Phase II, we see that low immunity and high permissivity result in a high environmental and total viral load. These viruses also do not go extinct, regardless of initial conditions. However, even a small increase in immunity leads to a relatively large decrease in viral load. As such, we associate this phase with “opportunistic” viral infections with high viral loads that only generally present in immune-suppressed and -compromised individuals, such as cytomegalovirus and JC virus [8, 42].

Moving forward, we therefore refer to the three phases as “acute” (Phase I), “opportunistic” (Phase II), and “chronic” (Phase III). We now go on to explore the dynamics of this model, beginning with the time to steady state, with two main questions in mind: (1) do the dynamics also reflect the phase transition, and (2) how are the dynamics of viral infections in the three phases related to the three classes of viral infections described above.

### Dynamics

#### Time to steady state

We now look at the time to steady state, shown in Figs. 7A, B (with convergence defined in Eqs. 9a,b). To determine a realistic timescale for iterations in our model, we consider experimental data from with real biological diseases.

**Fig 7.**
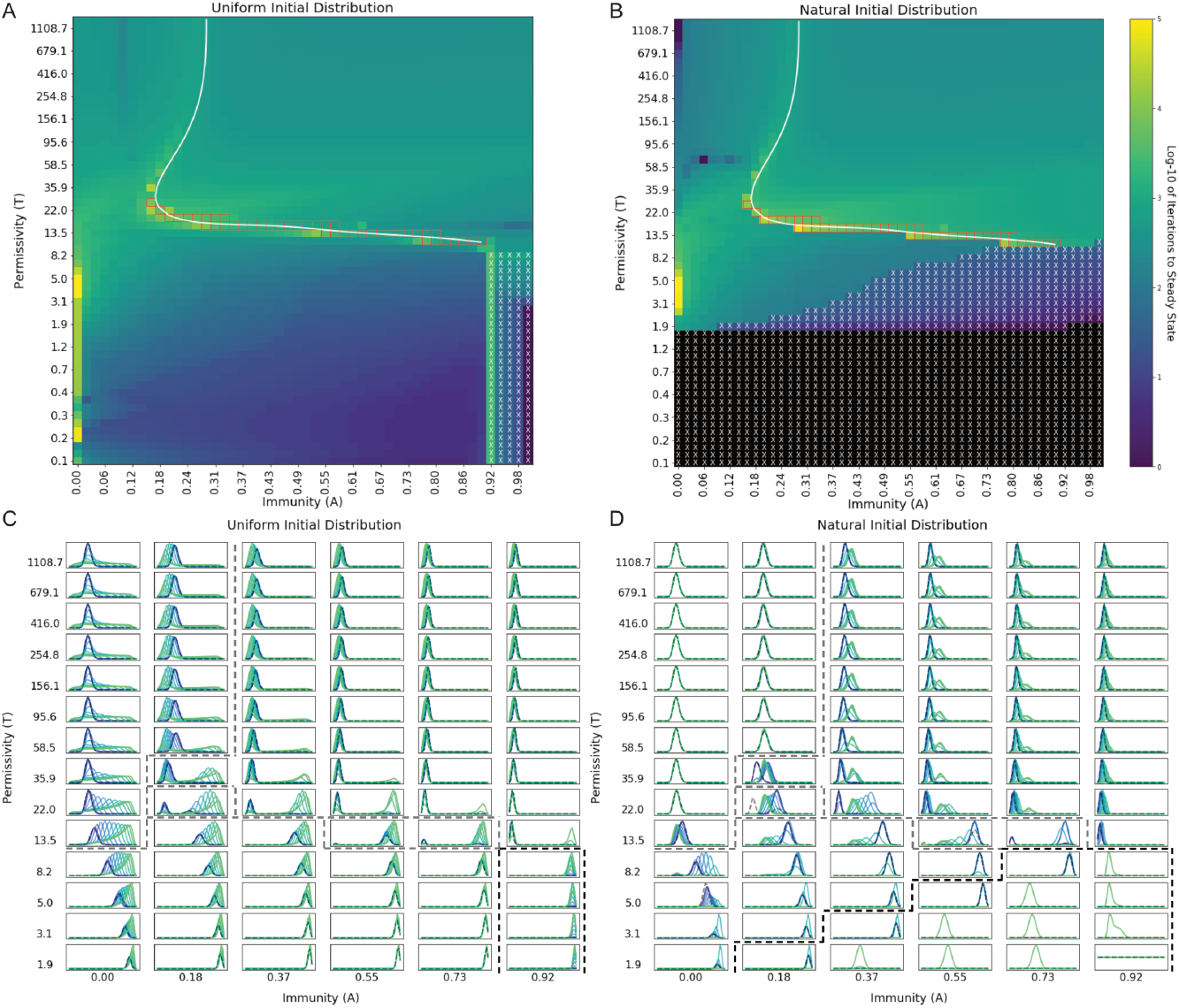
Convergence landscape is well-separated by phase boundaries and reflects a range of dynamic behaviors. (A) Iterations to convergence, Uniform initial distribution. (B) Iterations to convergence, Natural initial distribution. (C) Environmental match-number probability distribution scaled by viral load, uniform initial distribution. (D) Environmental match-number probability distribution scaled by viral load, Natural initial distribution. Panels A and B depict the iterations to steady-state on a log-10 scale as colors ranging from purple (1 iteration) to yellow (10^5^ iterations) and black (< 1 iteration). Panels C and D have an x-axis ranging from 0 to 50 and a y-axis scaled to show maximum dynamic range. Bimodal distributions are denoted by red boxes (A and B) or enclosure by gray dashed lines (C and D). Extinction is denoted by white X’s (A and B) or enclosure by black dashed lines (C and D).

Using infection dynamics data from studies of a range of viral diseases, including flu, and SARS-CoV-19, we can approximate the length of a full infection and replication cycle (i.e. cellular generation interval) as 0.5 days [43, 44]. Based on this approximation, we find that “acute” infections resolving in total viral extinction generally correspond to an illness duration of less than one month, similar to what is observed in the clinic [45, 46]. Further reinforcing this point is the observation that increasing immunity leads to more rapid viral clearance, as is to be expected for acute viral infections. In contrast, for “chronic” infections in the upper-right region of the phase diagram, the time to steady state ranges from hundreds to thousands of iterations. Using the approximate cellular generation interval for HIV-1 (2.6 days) for a single iteration in this region, we find that the system exhibits the months- to years-long timescale typical of chronic and persistent viral infections [47, 48]. For the “opportunistic” phase, we find times to steady state that are close to but shorter than those of the “chronic” region. These results give confidence that the model reflects biologically relevant dynamics and reinforces the association between different phases and different disease types.

We move on to other observations regarding Figs. 7A, B. In line with earlier results, we see that the order parameter-derived phase boundary (white spline) overlays neatly on the landscape of iterations to steady state, reinforcing the finding that nearly every property of this system reflects the phase transitions.

The most marked feature of this data occurs along the horizontal phase boundary, where we see a series of narrow horizontal regions exhibiting an unusually large number of iterations to convergence (yellow to yellow-green stripes). These long-lived simulations all exhibit bimodal match-number distributions, both at steady state (Fig. 4) and during the dynamics (Figs. 7C, D). We note that some points in these regions exhibit meta-stability (see S3 Fig) lasting many iterations (tens of thousands, in some cases) before finally and suddenly converging to steady state.

The metastable states explored around these values of *T* and *A* exhibit probability distributions and viral loads very close to those at steady state. We can therefore view the progression of these simulations as natural experiments in which the metastable states serve as initial conditions that are small perturbations from steady state. Seen in this way, the very long time to steady state serves as evidence of “critical slowing down,” a phenomenon typically seen near phase transitions in a wide variety of dynamical systems [49].

As mentioned, we also see, after large numbers of iterations, that metastable states abruptly shift to steady state. This phenomenon is known in the context of glassy dynamics as “quakes” [50]. Glassy dynamics typically involve a competition between closely-balanced environmental pressures (e.g., energy and entropy), reflecting a complex energy landscape with many similar minima that lead to long exploration times [51]. Such behavior is exactly what we see in our system (see, e.g., S3 Fig) for a subset of values of *T* and *A* adjacent to the horizontal phase transition. We also note that, although points in phase space along the horizontal boundary exhibit the long-lived behavior typical of competing pressures, the regions above and below that boundary show moderately to dramatically decreased iterations to steady state, reinforcing that the phase boundary is a narrow region of maximally competing viral adaptation strategies.

Examination of the dynamics provides additional insight into the causes of viral extinction. In Fig. 7B we see that the region of extinction consists of two blocks, shown in black and blue. At very low permissivity, viruses fail to survive beyond the first round of infection (black). At higher permissivity, viruses persist for relatively brief periods of time before going extinct (purple and blue, tens to hundreds of iterations). Thus, we can see that viral extinction in this model is a dynamic process that is strongly dependent on both the immunity and the initial conditions. Finally, we consider the time to steady state of viruses that survive in the “acute” region. At *A* = 0 and *T*≈ 3 − 5, we unexpectedly find an entire region of phase space in which the steady state of the system is oscillatory (bright yellow, S4 Fig). There, the system reaches a dynamic steady state with effectively infinite convergence time: no single steady-state quasispecies distribution is associated with those values of *T* and *A*. Instead, the system oscillates in viral load by up to 200 viruses and in mean match number by Δ*m* = 1 (more details in S4 Fig). Considered in the broader context of the “acute” region, we can interpret the periodic behavior as the dramatic culmination of a trend that appears throughout the entire region: the time to steady state is very low at high *A* and low *T*, and progressively increases as one diagonally approaches the subregion with oscillatory steady states. There, the pressures of permissivity and mutation (energy and entropy) are effectively equal, resulting in a finely balanced two-state system, moving slowly over time between one state and the other.

### Scaled distribution dynamics

We now consider the dynamics of the quasispecies distributions, shown in Figs. 7C, and D. The dynamics of the distributions give additional insight into two patterns in steady-state results seen earlier: 1) In the low *T* “acute” region, the peaks of steady-state distributions increase to higher *m* as *A* increases at fixed *T* ; 2) In the higher *T* “chronic” region, the peaks of steady-state distributions shift toward lower *m* as permissivity decreases at fixed *A*.

At very low *T*, in the “acute” phase, we see considerable mutation, as is characteristic for many viruses associated with acute infections. The pressure of permissivity dominates, preventing all but the highest match-number viruses from replicating. At lower *A* (*A*< 0.5), the viral quasispecies begin at high *m* but are able to shift over time to lower *m* by mutation. As immunity increases, however, viruses have less and less time to shift to lower *m* before being eliminated by the immunity, leading to steady-state distributions stuck at higher *m*. We note a similar effect as permissivity increases at low *A*, where the entropic pressure of mutation is increasingly able to push the virus towards *m* = 14, the point of highest viral sequence degeneracy (the number of different ways to form a virus of match number *m*).

In the higher-*T* “chronic” region, the pressure from immunity is greater than that of permissivity, enabling only low-*m* viruses to survive. The peaks of steady-state distributions in this region shift toward lower *m* as *T* is decreased at moderate *A* because, even with reduced infection and replicative capabilities, viruses are best able to survive by evading the immune response. The dynamic behavior in this region varies according to initial conditions because, while the “uniform” distribution already has low-*m* viruses, the “natural” initial distribution does not and thus needs to evolve viruses with that property. Notably, the “uniform” distribution simulations show very low-*m* viruses initially infecting, which then mutate and shift over time to increase *m* and reach their steady states. We see in particular near the horizontal phase boundary that, although higher-*m* viruses infect early on, they are cleared by the immune response, enabling the more evasive low-*m* viruses to remain in cells and replicate, albeit to a limited extent. This is an analogous effect to what we see in the “acute” region, where viruses with the other extreme value of *m* initially dominate and then slowly evolve into a less extreme steady state.

We also note that bimodality in the distribution dynamics is much more common than bimodality at steady state. For *T* and *A* around the horizontal phase transition (13.5 ≤ *T* ≤ 58.5, 0.18 ≤ *A* ≤ 0.73, we see the emergence of bimodality in the dynamics early on for the “uniform” initial distribution. However, over time, the dominant pressures for the relevant regions cause one of the two dynamic quasispecies to go extinct, except for at the phase boundary itself.

#### Viral load dynamics

We now discuss the dynamics of the viral loads inside and outside cells, which we show in Fig. 8. We observe two different timescales for the population dynamics: a faster “calibration” timescale that is strongly dependant on initial conditions, and a slower timescale reflecting the phase-specific viral strategy. This difference in behavior during shorter timescales reflects the effects of viral infection in humans, where early reactions to infections depend heavily on the individual, their immune response, and the viral load; while longer-term effects of a particular viral infection are more similar across larger populations.

**Fig 8.**
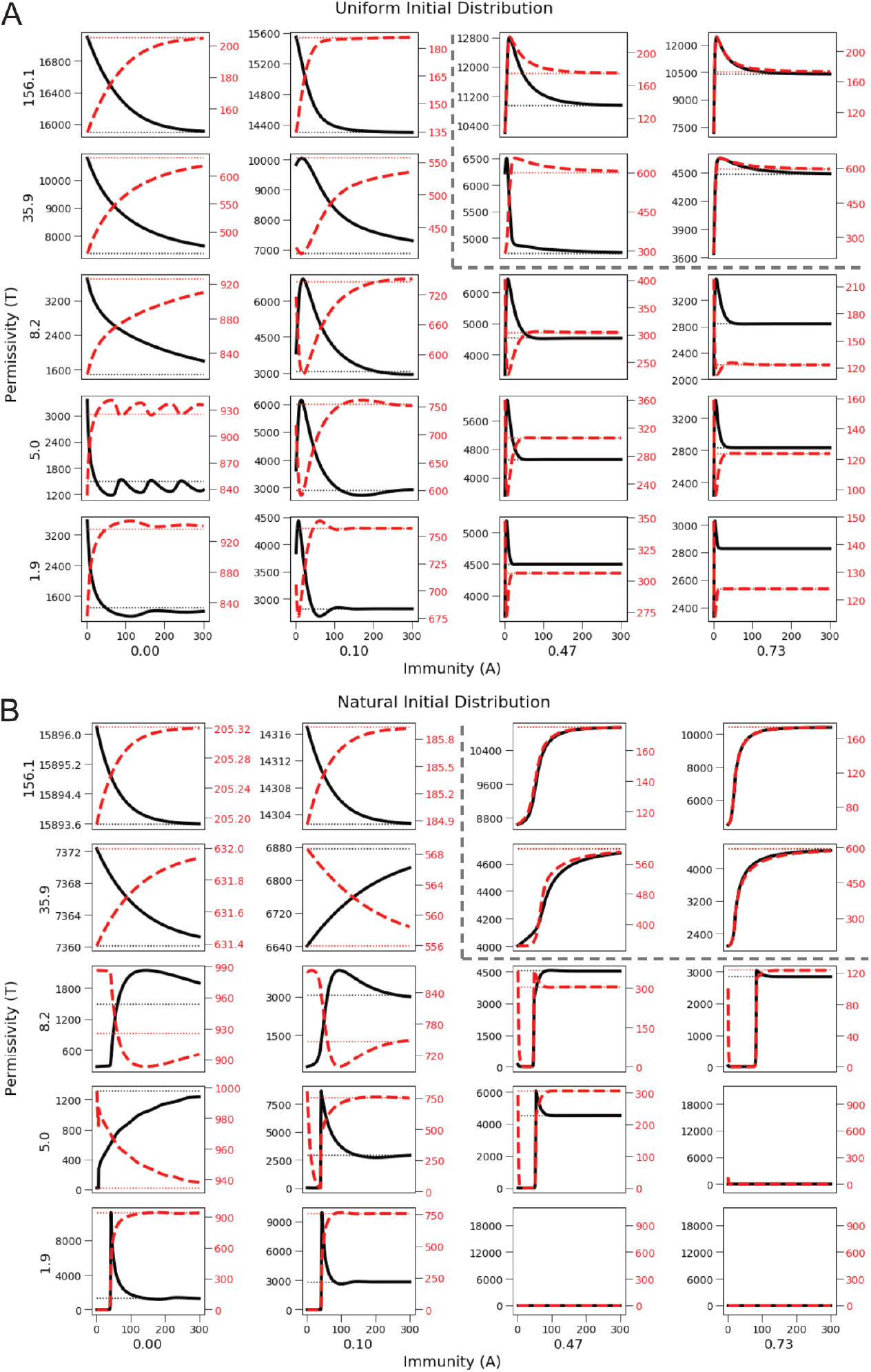
Dynamics of population size exhibit distinct behavior across the phase boundary. (A) Uniform initial distribution. (B) Natural initial distribution. Black axis and solid line, viruses in the environment. Red axis and dashed line, viruses inside cells. Black dotted line, steady-state value for viruses in the environment. Red dotted line, steady-state value for viruses inside cells. Horizontal axis is iteration number, a proxy for time. As in other figures, the phase boundary is overlaid as a dashed dark gray line between subplots, with points along the phase boundary boxed by the dashed line.

The overarching theme for the following results is that the phase boundary divides parameter space into regions of distinct relationships between the viral loads inside and outside cells for both initial conditions. We find two trends: the viral loads in the environment and within cells are either correlated or anti-correlated. In the “opportunistic” and “acute” regions, we observe anti-correlated behavior for both initial conditions. In these regions, the total viral load reaches its steady-state value during the fast timescale. Therefore, during the majority of the dynamics shown in Fig. 8, a change in the environmental viral load must lead to an equal and opposite change in the intracellular viral load.

By contrast, we observe correlated scaling of the viral loads in the “chronic” phase. In this phase, heightened immune clearance and lower match numbers slow the growth of the total viral load, which can only converge at the rate of its slowest constituent component. We note the remarkable scaling at high permissivity and high immunity, where the viral loads within cells and in the environment grow in lockstep, in particular for the natural initial condition (Fig. 8B). Importantly, the viral load dynamics in this phase show how a high total viral load can be reached in non-immunocompromised humans.

Finally, we note that it is not universally true that the within-cell and environmental viral loads grow monotonically throughout disease progression. Rather, we see a complex set of dynamics, where the viral loads in the environment and in cells have different values and strive through the various pressures to meet the most fit state relative to the given environmental conditions. Therefore, while it is possible to make general statements regarding the growth of the system from its initial conditions, the fine details of change depend heavily on the relative interplay of the five systemic pressures. The whole of this serves as a prediction for the infection dynamics of viral diseases, for which time-resolved *in vivo* observations are not yet possible.

## Discussion

In this work, we have shown how a collection of viruses that infect cells, face an immune response, replicate, and mutate, can lead to a system exhibiting multiple behaviors corresponding to three distinct phases of a phase diagram separated by both first-order and continuous phase transitions. These phases emerge in a space we construct from the two main parameters of our model, permissivity (*T*) and immunity (*A*), and exhibit self-consistent properties unique to each phase.

Collectively, the pressures of permissivity, immunity, mutation, entropy, and the available pool of infectable cells compete, leading to three distinct classes of viral populations that correspond to the three different phases. In the high-permissivity and low immunity phase (Phase II), viruses are primarily directed to steady state by entropy in the form of mutational burden. In the high permissivity and moderate-to-high immunity phase (Phase III), viruses are driven to steady state by an attempt to evade the immune response (through decreased *m*), enabled by the uniformly high infection rate at high permissivity. In the low permissivity phase (Phase I), infection and replication are only possible at higher match number, leading to an immunity-dependent push toward steady-state quasispecies distributions with decreasing width and increasing match number. Taken to the extreme, the restrictions of low permissivity and high immunity lead to the complete extinction of viruses, even for the uniform initial distribution.

We link the three general classes of viruses from our model to infection types affecting humans in real life: acute, chronic, and opportunistic. Firstly, we suggest that the class of viruses below the horizontal phase boundary (Phase I), due to their persistent replicative ability at nearly all levels of immunity, corresponds to “acute” viral infections brought on by viruses that are highly adaptable and generally infectious, such as the influenza viruses, rhinoviruses, and many coronaviruses [36, 37, 52, 53]. Secondly, viruses at a broad range of immunity above the horizontal phase boundary (Phase III) can be identified with “chronic” infections: low match numbers enable them to persistently evade the immune response. Such behavior matches well onto the set of viruses that stay primed in the body for months to years, such as varicella-zoster (chicken pox/shingles), hepatitis B and C, and HSV [38–41]. Lastly, at high permissivity and low immunity (Phase II), viruses exhibit moderately low match numbers, corresponding to highest viral sequence degeneracy. These viruses map well onto “opportunistic” viral infections that only generally present in immune-suppressed and -compromised individuals, including cytomegalovirus and JC virus [8, 42].

Our results reinforce this interpretation in several ways. Firstly, in the “chronic” phase, we find that there is no clear resolution of viral infection even if immunity is increased, consistent with the physiology of long-term, chronic diseases. We contrast that with the “opportunistic” phase, where even a small increase in immunity leads to a relatively large decrease in viral load, as is to be expected for this type of disease. Finally, in the “acute” phase, the level of immunity has a large effect on the rate of recovery, as evidenced by the corresponding decrease in time to steady state and increase in extinction probability. This reflects the typical resolution of acute viral infection after the development of an immune response.

We can also ask what the model would suggest to drive viruses causing these diseases to extinction. Firstly, we judge it more likely that an individual receives an initial bolus of viruses that more closely resembles the “natural” distribution than the “uniform” distribution. For the “natural” distribution, there is a large region of viral extinction in our phase space. With this in mind, we can now analyze infection types. For an “acute” infection, the only guaranteed paths in our model to viral extinction are to decrease permissivity and/or increase immunity. However, in those patients for whom an increase in immunity is not possible, our model suggests that only a drastic decrease in *T*, likely via inhibiting general aspects of viral replication, will significantly decrease viral load. For the other two types of infections, only by decreasing *T* to meet the horizontal phase boundary can full extinction be attained. Supporting that idea, we note that for patients with “chronic” diseases such as hepatitis C, increasing the level of immunity is often insufficient on its own to cure the underlying disease, a result also found in our model [54, 55].

For a broader initial distribution of viruses, our model leads to a non-zero viral load in steady state for all but the highest levels of immunity. Based on our model, we posit that wellness may be attained without the total clearance of viruses. While high levels of viral load more often correspond to severe disease, low viral loads may not lead to any symptoms; indeed, prior work has shown that a non-negligible proportion of the population tested positive for flu antigen but did not exhibit any symptoms [56, 57].

In considering future work to improve the model, we suggest four high impact modifications corresponding to increasing levels of complexity. Firstly, rather than keeping secondary parameters constant, such as the mutation rate and fecundity, we believe that it would be useful to see the effects of varying individual parameters across different phases and within individual simulations, reflecting the variation in mutational and replicative rates across different viruses [58, 59].

Secondly, we propose implementing an immune module that is capable of actively updating and expanding its memory. In its present form, our model’s immune response exhibits a static memory with a solely match number-dependent level of response representing a fixed past history. In humans, however, the adaptive immune system actively recognizes new threats, generates a response, and stores the capability to respond in memory [60]. In its simplest form, adaptive immunity could be modeled by dynamically changing *A* during a single simulation. A truly adaptive immune system would likely substantially increase the size of extinction regions, as static quasispecies would be hard pressed to exist for extended periods of time.

Thirdly, it is also possible to extend this model beyond the cell-virus regime to include a broader environment, allowing for interactions between individuals or populations, as is typical in standard epidemiological approaches. One way to include patient-patient interactions would be to suddenly introduce new viral quasispecies during the course of a simulation. Alternatively, subsets of cells could be assigned distinct receptor sequences and then allowed to infect one another.

Finally, we discuss modifications that can be made to inform the development of novel viral therapeutics by directly incorporating and evaluating the effects of existing antiviral therapeutics into simulations. Doing so requires a moderate level of modification to the existing code, as antivirals interrupt viral replication across a wide range of points throughout the infection cycle; however, this modification would effectively demonstrate the usefulness of this model in evaluating novel therapeutics at an early stage for low cost.

We believe that aspects of our model may be tested through biological experimentation in mice or other model organisms. Although our model cannot at this time provide quantitative predictions for experimentally measurable quantities, it should still be possible to test qualitative aspects. In particular, we are deeply curious about testing for the existence of the reported first-order phase transition in a biological system. We describe one possible set of experiments in the appendix (S1 Appendix, section 7).

## Conclusion

In summary, we find that our relatively simple analytic model for within-host viral infection and evolution produces a wide variety of lifelike features and interesting physical phenomena. Results for steady states and the dynamics reveal an underlying set of phase transitions and crossover regions. The phase transitions and crossover regions partition three main phases corresponding to distinct classes of quasispecies behavior, both at steady state and in the dynamics, which we map onto three broad categories of human-infecting viruses. We observe a wide range of phenomena across multiple timescales, including glassy dynamics, bimodal quasispecies distributions, and periodic steady states. Based on these results, as well as the promise of extensions made possible by its modular structure, we believe that our model can be used to gain deeper insight into the complex interactions between infection mechanisms, host immune defenses, and the dynamics of viral evolution.

## Supporting information

**S1 Fig. Perturbation expansion validates quadratic approximation**.

**S2 Fig. Perturbation theory accurately predicts low-permissivity population sizes**.

**S3 Fig. Metastability of slowly-converging points in phase space**.

**S4 Fig. Periodic steady state at** *A* = 0,*T* = 5.05.

**S1 Appendix. Supporting information**. The appendix contains additional text, derivations, and figures for the following sections: Section 1 - Infection rate derivation; Section 2 - Mutation matrix description; Section 3 - Linear dependence of total viral load on immunity in Phase I; Section 4 - Perturbation theory expansion for low-permissivity results; Section 5 - Metastability results; Section 6 - Periodic orbits; Section 7 - Suggestions for experimental evaluation of model.

## Acknowledgments

We acknowledge support from the Center for Cellular Construction (NSF DBI-1548297), and from NIH grant R35 GM130327 (WFM). WFM is a Chan Zuckerberg Biohub investigator. This work was performed at the Aspen Center for Physics, which is supported by National Science Foundation grant PHY-1607611 (BAJ). This material is based upon work supported by the National Science Foundation Graduate Research Fellowship under Grant No. 1650113 and by the National Institute of General Medical Sciences of the National Institutes of Health under award number T32GM008284 (GRL).

## Supporting Information for

## 1 Infection rate

In this section we derive equations for the infection rate in our model. To start, we will calculate the probability that a single virus will infect any one single cell of the *c* total cells encountered within the host. We do this because, in this framework, each virus will either infect a single cell or not infect at all. It may be the case that some proportion of cells are already occupied, which will reduce the probability that a virus infects a cell. The probability of a single virus infecting is

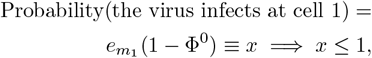

where 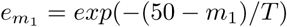 (Eq. 1 in main text) is the probability that a virus of match number *m*_1_ can infect a single isolated cell at permissivity *T* and Φ^0^ is the probability that the cell has already been infected (the sum over all m *m* of the probability that the cell was already in an occupied state by a virus of match number *m*).

We can now calculate the probability that the virus infects any one cell in the pool. This value is equal the sum of the probabilities that: the virus lands and infects at the first cell (cell 1); that it does not infect at cell 1, but does at cell 2; that it does not infect at cells 1 or 2, but does at cell 3; …infects at none of the previous cells, but does so at cell *c*. Mathematically, this is represented by:

Probability(the virus infects the second cell but not at the first cell)

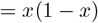

Probability(the virus infects the third cell but not at the first two cells)

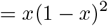

note that this is equal to the quantity *x*(1 − *x*(1 − *x*))

Probability(the virus infects the fourth cell but not at the first three cells)

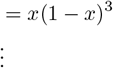

Probability(the virus infects the c-th cell but not at the first c-1 cells)

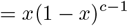

Probability(the virus infects any cell)

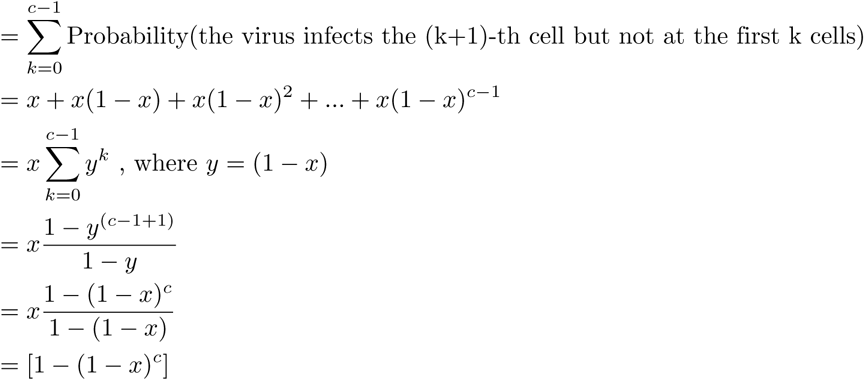

Here we can see that the probability for a virus to infect at any one cell is equal to one minus the probability that it does not infect at any cell. We note that the spatial distribution of the cells has disappeared from these expressions, consistent with model. Substituting in the original variables gives

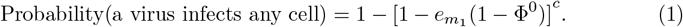

How do we account for the distribution of viruses? The probability that a virus emerging from the collection of viruses in the environment of being match number *m* is equal to *P*_*m*_. Once the virus has begun attempting to infect cells, the distribution has no further bearing on that particular virus’s ability to infect, as it already has a match number *m*. Therefore, the probability that the first virus infects with match number *m* is

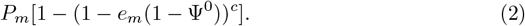

To convert this probability of the first virus infecting a particular cell into one of infecting any of the cells in the infectible pool, we divide by *c* to obtain the total additional infection probability added by the first virus, 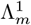:

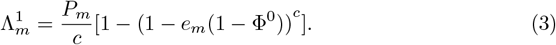

The total probability per-cell, including the previously occupied cells, is now updated to be:

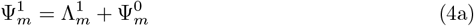

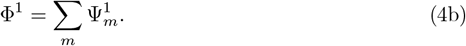

In our system, the initial cellular occupation is given by the viruses remaining in cells, Ψ^*R*^:

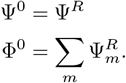

Once the first virus has had an opportunity to infect, the second virus emerges and attempts to infect. The existing level of cellular occupation that it encounters must now take into account both the pre-infection probability of occupation and the probability that the first virus successfully infected:

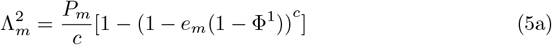

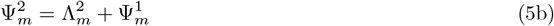

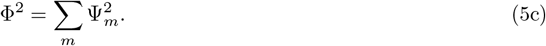

This process continues until the *N*-th virus has had an attempt to infect:

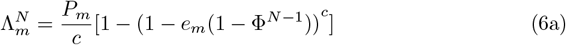

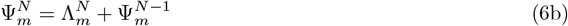

Thus, for a system with *N* viruses, the distribution of viruses that have infected cells is

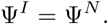

We note that the number of viruses in the environment after reproduction and mutation is a real number. However, Eq. 6b requires an integer number of viruses. Therefore, to determine the value of Λ for real *N*, we linearly interpolate the value of Λ between the two whole numbers of viruses nearest to *N*:

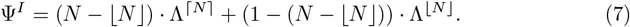

## 2 Mutation Matrix

Viral progeny may have match numbers that differ from their parent virion by Δ*m* = − 1, 0, +1. The equations determining the probabilities of mutation were derived previously [1] and are repeated here:

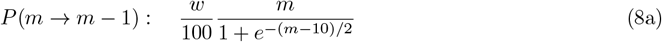

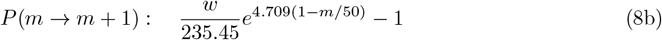

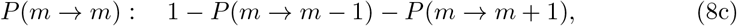

where *m* is the match number and *w* is the degeneracy of the chosen host cell receptor sequence (with value 0.7867 from previous work).

## 3 Derivation of linear dependence of total viral load on immunity in Phase I

For nearly all simulations in this regime, the total infection probability at steady state is approximately 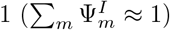. Also, the match-number distributions at steady-state are centered and concentrated at relatively high *m* (*m*> 30), for which the immune response is effectively constant and equal to *A*. Therefore,

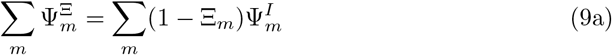

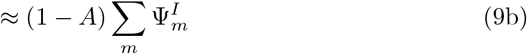

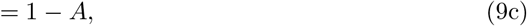

demonstrating permittivity-independent linear scaling of total viral load with immunity.

Within the same phase, the viral load inside cells contrasts with that in the environment by showing a super-linear and monotonic decrease from high to low value as immunity increases. This can be explained as follows. As immunity increases, the match number distributions shift towards higher *m*. At such low permittivity, the probability that viruses remain inside cells (1 - *e*_*m*_) exhibits exponential decay as a function of increasing *m*. Therefore, as immunity increases, the probability that viruses remain inside cells decreases super-linearly.

This result provides another explanation for the scaling behavior of the viral load in the environment: subtracting a super-linearly decaying function from a linearly decaying function gives a maximum in the difference.

## 4 Low-permissivity perturbation theory derivation and results

In order to understand the limiting behavior of the model, we applied perturbation theory appropriate for the low-permissivity regime to derive steady-state solutions for the within-cell and environmental population sizes at steady state.

At sufficiently low permissivity, only viruses with *m* = 49 and *m* = 50 should make non-negligible contributions to the system, so we constrain our consideration to include only those types of viruses. Also, given such low permissivity, we expect the replicative ability of *m* = 49 viruses to be negligible. In order to stay in the perturbative regime, we must remain at sufficiently low permissivity such that *m* = 49 viruses do not reproduce, as their reproduction would lead to an expansion of the variety of viruses present in this system (thereby contradicting our initial assumption requirement for only two types of viruses). Additionally, any viruses that remain inside cells after reproduction must have match number equal to *m* = 49, as *m* = 50 viruses always reproduce with probability 1 and escape into the environment: thus, the entire within-cells viral population is exactly the population of *m* = 49 viruses inside of cells. Therefore, whenever viruses replicate into the environment, they must come from *m* = 50 progenitors in a proportion determined by the mutation matrix: *P* (50 → 49) ≈ 0.39 and *P* (50 → 50) ≈ 0.61.

To derive steady state conditions at any immunity, we assume that the system is near steady state and has some population inside cells (after reproduction) equal to *μ*. During the next iteration, just after infection by viruses in the environment, the cells will be fully occupied if there were sufficiently many infecting viruses with *m* = 50 (*m* = 50 viruses have a near-certain probability of infecting unoccupied cells).

To calculate the updated infection probability of the *m* = 49 and *m* = 50 viruses, we need to know the ratio of their infective abilities and also have some knowledge of their infective abilities as a function of the prior level of infection, *μ*.

We first note that, when *μ* = 1, there is no additional infection. With this in mind, we start by considering the additional infection of *m* = 49 and *m* = 50 viruses to each be proportional to (1 *- μ*) with different coefficients.

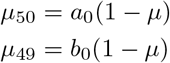

where *a*_0_ >> *b*_0_.

Note, however, that these equations would imply that the infectibility of *m* = 50 and *m* = 49 have a constant ratio for any initial level of infection *μ*. We know this is not correct, because for *μ* close to 1, the ratio of *μ*_49_*/μ*_50_ = *b*_0_*/a*_0_ must be much less than its value when *μ* = 0. Given that new infection by *m* = 49 viruses is more sensitive to the prior level of infection than that by *m* = 50 viruses (Fig. 2A of main text), we now try the next-simplest approximation and say that:

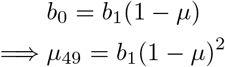

We now impose the sum rule, which says that the new total infection by *m* = 49 and *m* = 50 viruses is equal to 1:

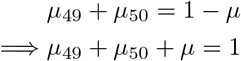

Thus,

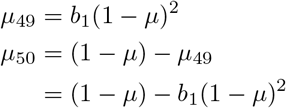

To test whether this is a reasonable approximation for infection at very low permissivity, we need to choose a permissivity satisfying the aforementioned constraints but is also sufficiently large that simulations at that permissivity will reach steady-state in a reasonable amount of time. Accordingly, we choose a sample permissivity of *T* = 0.1, as *e*_49_ ≈ 4.5 * 10^−5^.

To test this, we calculate the cell occupancy just after infection for *m* = 49 and *m* = 50 viruses. In particular, we assume 1000 cells and 10,000 viruses (however, the calculation is insensitive so long as the number of viruses exceeds 3,000 for *c* = 1000). The output of the infection process gives

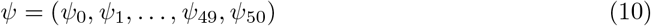

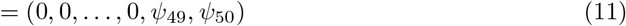

with our hypotheses:

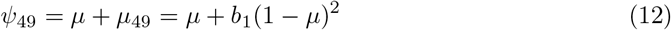

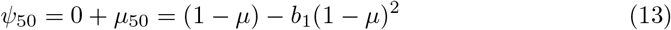

We then isolate *b*_1_(1 − *μ*) from each of the hypotheses and compare them to the calculated results:

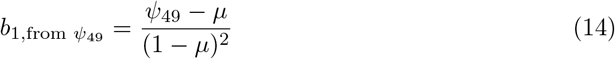

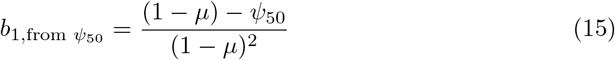

In Supplementary Figure 1, we plot *b*_1_ as determined from each of *ψ*_49_ and *ψ*_50_, as well as the average value of *b*_1_ calculated across all initial *μ*. We find that *b*_1_ is very nearly constant across all initial *μ*, that the values of *b*_1_ are identical regardless of whether they are calculated from *ψ*_49_ or *ψ*_50_, and that the averages across all initial *μ* are the same for both *ψ*_49_ and *ψ*_50_. This validates our initial approximation.

Given this result, we can now determine the self-consistent, steady-state solutions for the viral populations. We take *b*_1_ to be the value calculated for *μ* = 0 so as to be consistent with the perturbative expansion.

We begin with some prior infection level *μ*. As determined above, the cell occupancy just after infection is:

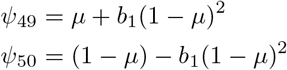

where *b*_1,*μ*=0_ = 0.01423388 for *T* = 0.1.

Immune activity reduces the viral population as expected:

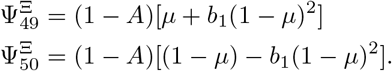

The viral load within cells after reproduction is:

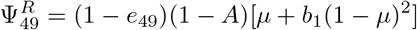

and, as *e*_49_ ≈ 10^−5^ << *b*_1_ ⇒ 1 − *e*_49_ ≈ 1,

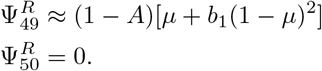

As only *m* = 49 viruses will remain inside cells, the final within-cell viral population is:

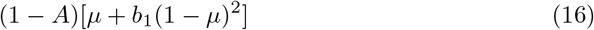

To determine the steady-state values, we now set the final within-cell population to *μ*, giving us a final, self-consistent equation for the within-cell population at steady state in the perturbative regime:

**Supplementary Fig 1.**
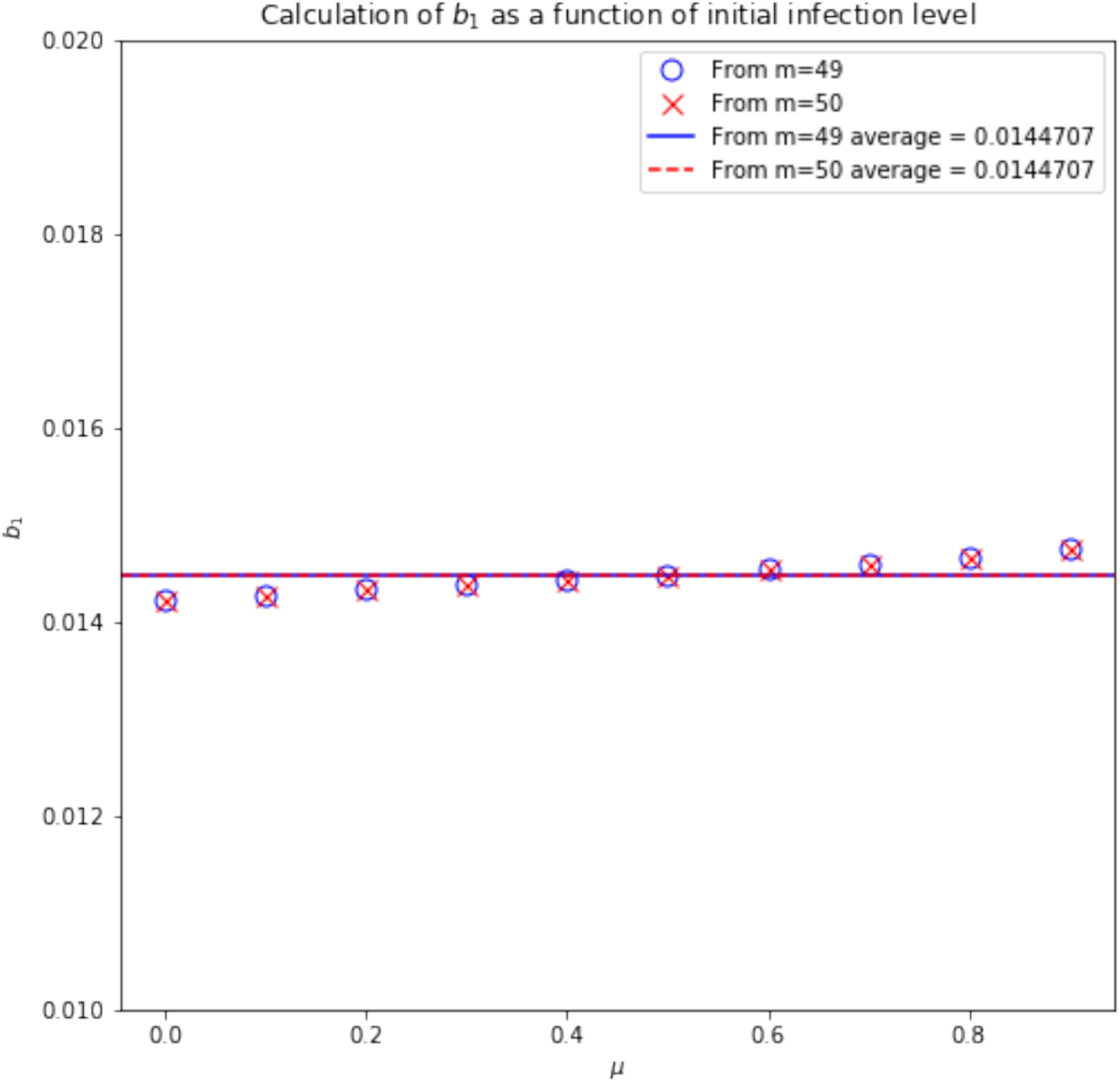
Perturbation expansion validates quadratic approximation. Blue solid line: average of *b*_1_ as calculated from *ψ*_49_. Red dashed line: average of *b*_1_ as calculated from *ψ*_50_. Blue O’s: values calculated from *m* = 49 expression. Red X’s: values calculated from *m* = 50 expression.

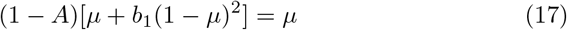

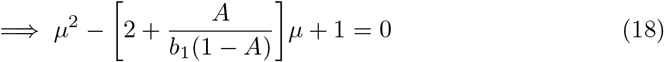

This is a standard quadratic equation in *μ* with solutions:

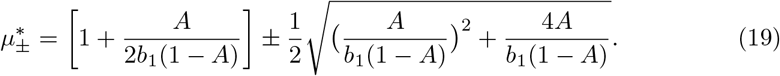

Inspection of the roots shows that the ‘+’ root exceeds 1, while the ‘−’ root falls between 0 and 1, so we use the ‘− ‘ root moving forwards.

Given that the predicted within-cell population at steady-state is equal to *μ*\* (*A*), the corresponding predicted environmental population at steady-state (scaled to 0-1) is:

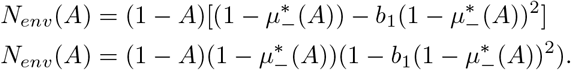

With these results in hand, we compare our predictions to the data from our simulations, as shown in Supplementary Fig. 2. We see that there is excellent agreement between the predictions and the data, except for where the simulations go extinct. Relative percentage errors do not exceed 1% for either computed quantity.

The figures demonstrate that our approximation, assuming that cells are fully occupied after infection, deviates from the data at both *A* = 0 and *A* ≈ 1. At *A* = 0, further analysis of the simulation dynamics shows that the system continues to evolve very slowly towards a state of full cell occupancy, at which point no further viruses from the environment can infect and reproduce, causing the environmental viral load to go to 0. At *A* ≈ 1, study of the infection equations show that complete filling after infection breaks down when *N* ≈ 2*c*. Translating this into the self-consistent steady-state equations gives a critical *A*\* at which this happens:

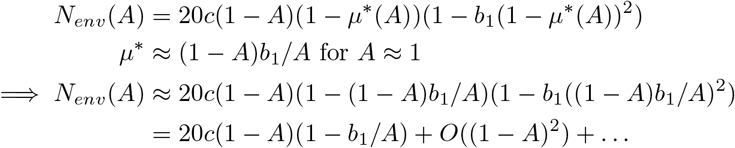

When 2*c* = *N*:

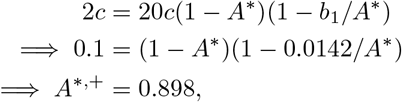

consistent with the data.

**Supplementary Fig 2.**
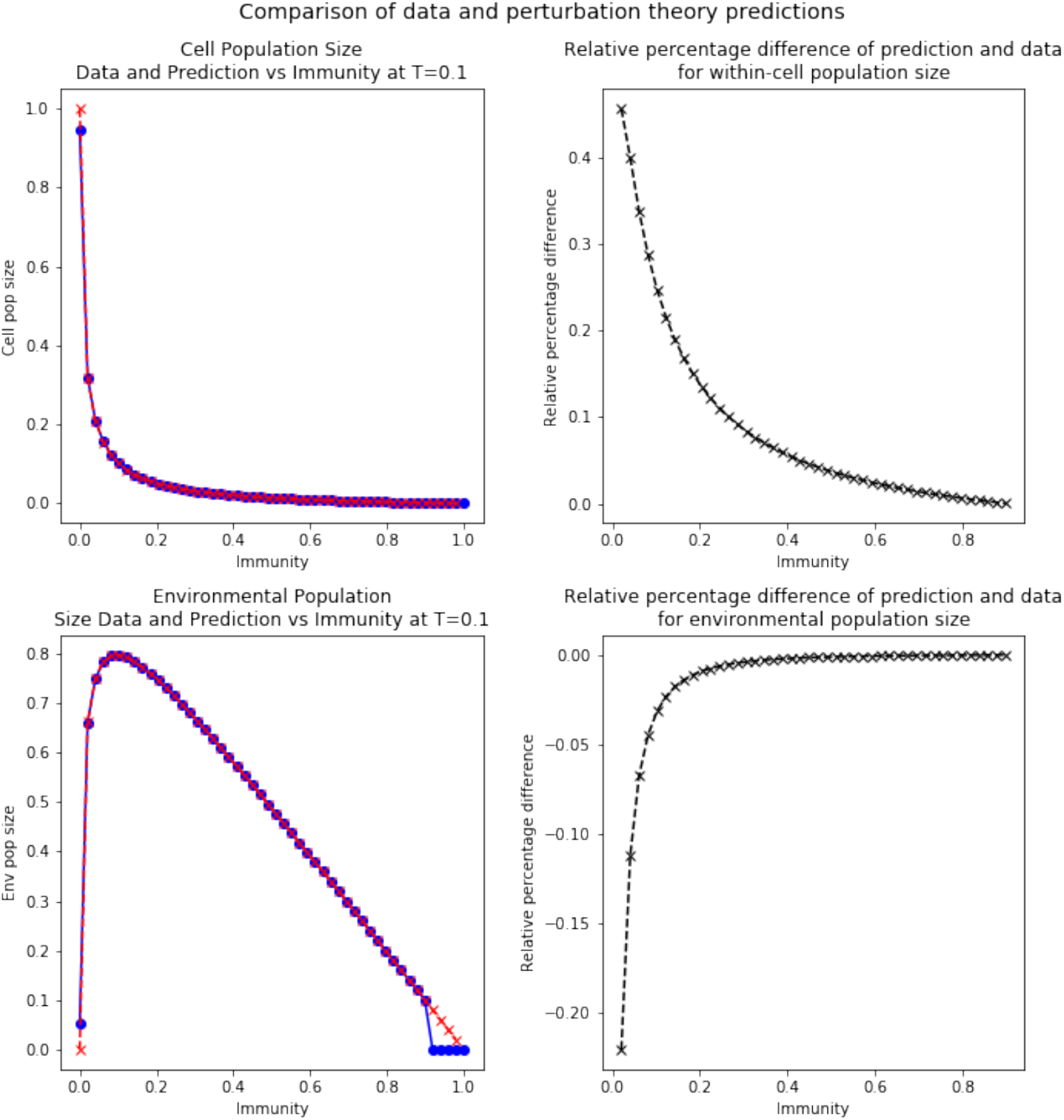
Perturbation theory accurately predicts low-permissivity population sizes. (A) Within-cell population size. (B) Relative percentage error of prediction and data for within-cell population size. (C) Environmental population size. (D) Relative percentage error of prediction and data for environmental population size. Red X’s: predictions from perturbation theory. Blue O’s: values from uniform initial distribution simulation, T=0.1.

## 5 Metastability of slowly-converging points in phase space

In Supplementary Fig. 3 we show evidence of the system reaching metastability that lasts for thousands of iterations. Please note the large scale of the iteration axes. Additional tests for stability (simulation out to one million iterations) has show that the system stays at the apparent final state to accessible limits.

**Supplementary Fig 3.**
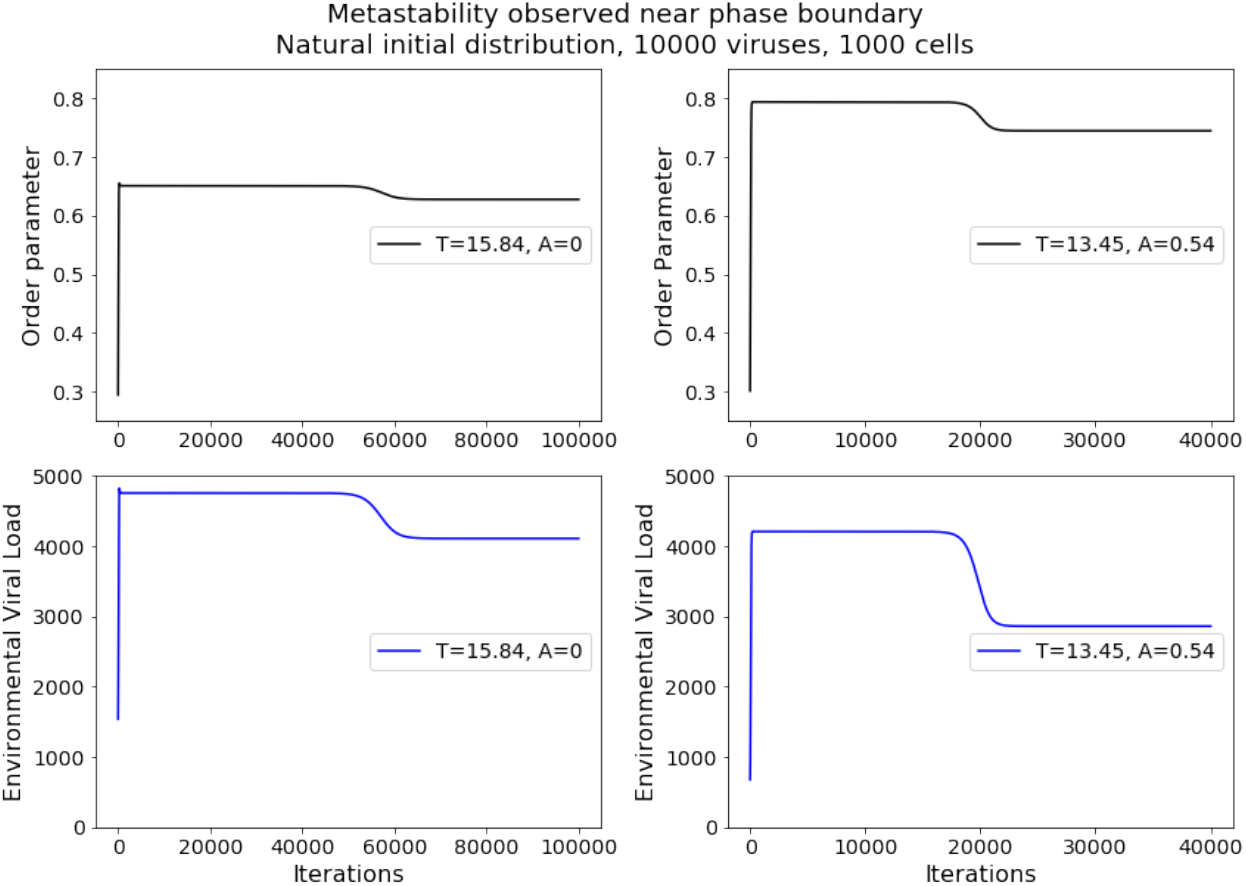
Points of very long convergence time demonstrate metastability persisting for thousands to tens of thousands of iterations.

## 6 Periodic orbits

At zero immunity and *T* between 3 and 6, unexpectedly we find for both initial conditions oscillatory-like behavior, which converges to periodic or pseudo-periodic behavior. In Supplementary Fig. 4 we show, for *T* = 5.05, that both the order parameter and population size oscillate, with the order parameter varying by 2% and the population size varying by 13%. The values of these parameters exhibit periods of ≈ 8000 iterations and continue to do so across the majority of the simulation. The magnitude of these variations, especially compared to the number of significant figures we keep for our calculations, suggests to us that the results are true reflections of the model’s behavior and not an unintentional result of rounding or other numerical phenomena.

Additionally, we find that the observed orbit is stable. In order to test the stability, we induce large perturbations towards the end of simulations to try to send the simulation toward a different steady state (perturbations include: within cell match number distribution, environmental match number distribution, and environmental viral load). In each case and in combination, all perturbations lead to a return to the same steady-state oscillatory results.

**Supplementary Fig 4.**
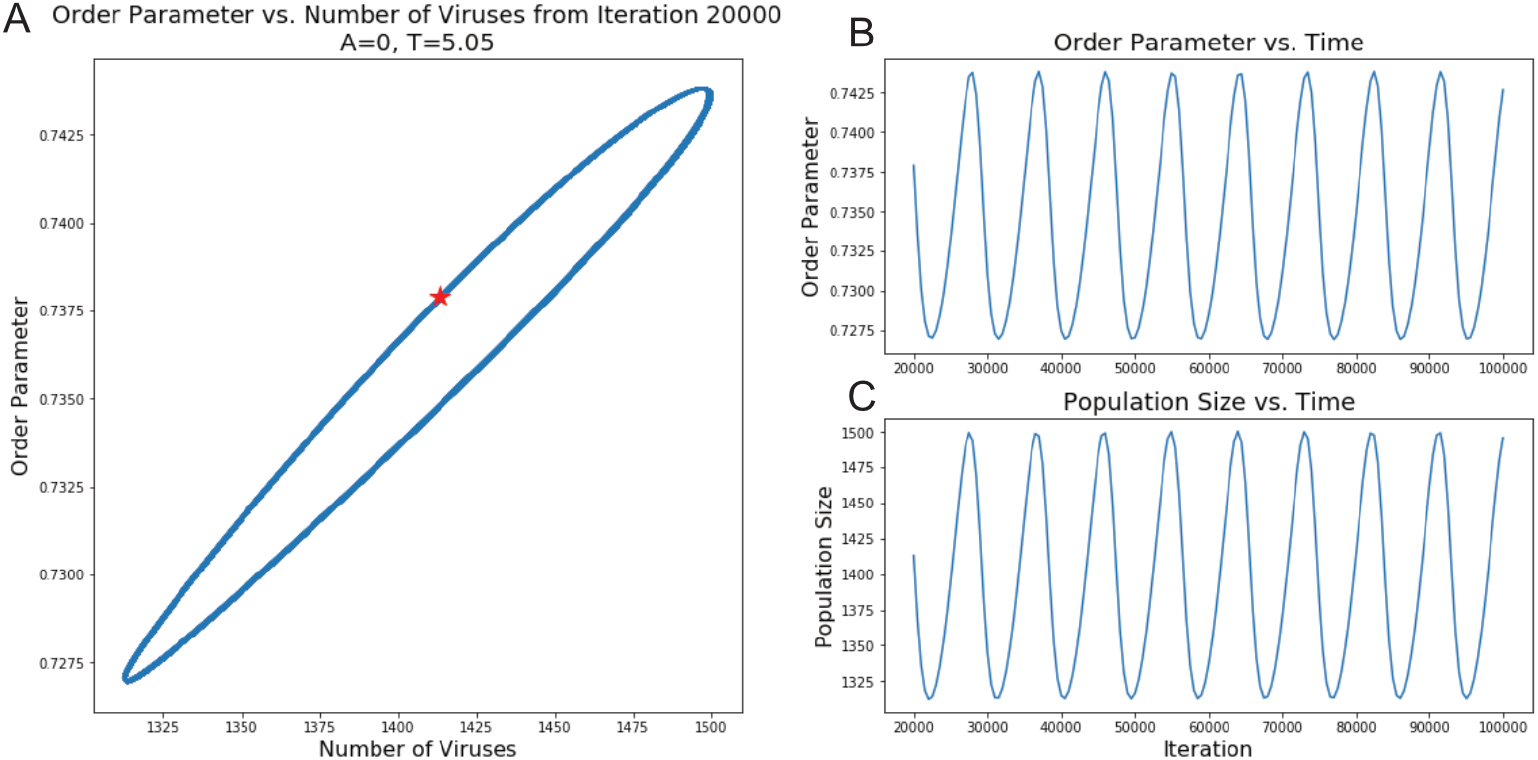
Stable periodic orbit appears at low permittivity and zero immunity. (A) Order parameter and viral population size exhibit periodic behavior for tens of thousands of iterations. Red star indicates position at iteration 20000. (B) Order parameter value of viruses in the environment from iteration 20000 onward. (C) Population size of viruses in the environment from iteration 20000 onward. Data in (B) and (C) are sub-sampled every ≈ 500 iterations for visual clarity.

## 7 Experimental testing of model

As described in the Discussion, we believe that it would be interesting to evaluate this model in an experimental context to gain insight into the relationships between pathogen population dynamics, host defenses, and other pressures. We envision a minimal viable test as follows. First, given that the first-order phase transition is most apparent at values of *A*> 0.3, the host should demonstrate a moderate but suppressed level of immune response. Next, a range of different strains of the host need to be generated to correspond to a wide range of permissivity; this can be done either by systematically modulating either the selectivity or number of the main host receptor engaged by the virus for cellular entry. Lastly, for each host strain, a large number of infection cycles need to be carried out, with fresh host cells of the appropriate strain replenished after each cycle, potentially using a chemostat or continuous-flow technique. If an analogous first-order phase transition is present in the experimental system, it should appear in the form of a sharp extinction of one viral quasispecies, followed by the emergence of a region favorable to a drastically different viral type on the other side of the boundary, with the possibility of a limited amount of simultaneous presence in the transition region. A complementary experiment would involve rapidly shifting a steady-state viral quasispecies from a high permissivity host strain to a low permissivity strain, which we predict will lead to rapid viral extinction. To avoid the complexities of mammalian viral-host interactions, the aforementioned experiments may be able to be performed in a suitable bacteriophage-bacteria system. If that is the case, it would be interesting to see if the transition in quasispecies type corresponds to a transition between lytic and lysogenic phage behaviors.

